# Enhanced GPP synthesis by Erg20p-peptide fusions and biomolecular condensates boosts monoterpene production in yeast

**DOI:** 10.64898/2026.04.28.721314

**Authors:** Ke Xu, Archontoula Giannakopoulou, Virginia Jiang, Saurabh Malani, Mackenzie T. Walls, Clifford P. Brangwynne, José L. Avalos

## Abstract

Monoterpenes are a diverse class of natural products with broad industrial and pharmaceutical applications. While there is great interest in transitioning their production from chemical synthesis and natural source extraction to yeast bioprocesses, this approach remains limited by the dual functionality of the endogenous farnesyl diphosphate synthase Erg20p, which produces the monoterpene precursor geranyl diphosphate (GPP) but favors its subsequent conversion to farnesyl diphosphate (FPP). To address this limitation, we recruited Erg20p and monoterpene synthases into synthetic membraneless organelles, improving production. In doing so, we found that short C-terminal peptide fusions used for recruitment also significantly enhance GPP synthase activity relative to FPP synthase activity. The combined effects of metabolic spatial organization and GPP synthase activity enhancement significantly boost production of different monoterpenes, including geraniol titers exceeding 4 g/L. The strategies presented here can be readily integrated with other traditional metabolic engineering approaches to build yeast strains with high levels of monoterpene production.

## Introduction

Monoterpenes, including monoterpene alcohols (e.g., geraniol and linalool) and esters (e.g., geranyl acetate and linalyl acetate), are widely used in the flavor and fragrance industries for their pleasant aroma^1,2^. Additionally, some monoterpenes exhibit a broad range of pharmacological properties, including anti-inflammatory, antimicrobial, and antinociceptive activities. Current industrial monoterpene production relies primarily on extraction from plant sources and chemical synthesis, both of which present sustainability challenges. Plant extraction methods are resource intensive (requirements include land, water, fertilizer, and pesticides), generate significant waste streams and availability is highly dependent on weather conditions and climate^3^. Chemical synthesis methods, on the other hand, depend on non-renewable petrochemical feedstocks^4^. Furthermore, growing global demand for monoterpenes is placing increasing pressure on these systems^5^. Consequently, biosynthesis of monoterpenes using engineered microorganisms and sustainable carbon sources has emerged as an attractive alternative to address these sustainability challenges and meet the growing demand^6,7^.

In the monoterpene biosynthetic pathway, two 5-carbon precursors, isopentenyl diphosphate (IPP) and dimethylallyl diphosphate (DMAPP), synthesized via the mevalonate (MVA) pathway, are condensed by the GPP synthase Erg20p to form geranyl diphosphate (GPP, C_10_), the key precursor for monoterpene biosynthesis (Figure 1a)^8^. The same enzyme also functions as a FPP synthase, catalyzing the subsequent condensation of GPP with IPP to form farnesyl diphosphate (FPP, C_15_). The Erg20p active site contains an allylic (A1) site, which binds DMAPP or GPP, and an homoallylic (HA1) site, which binds IPP. Erg20p undergoes homodimerization, a configuration essential for its catalytic activity, and its dimerization state has been shown to cooperatively regulate terpene biosynthetic flux^9^.

**Figure 1.**
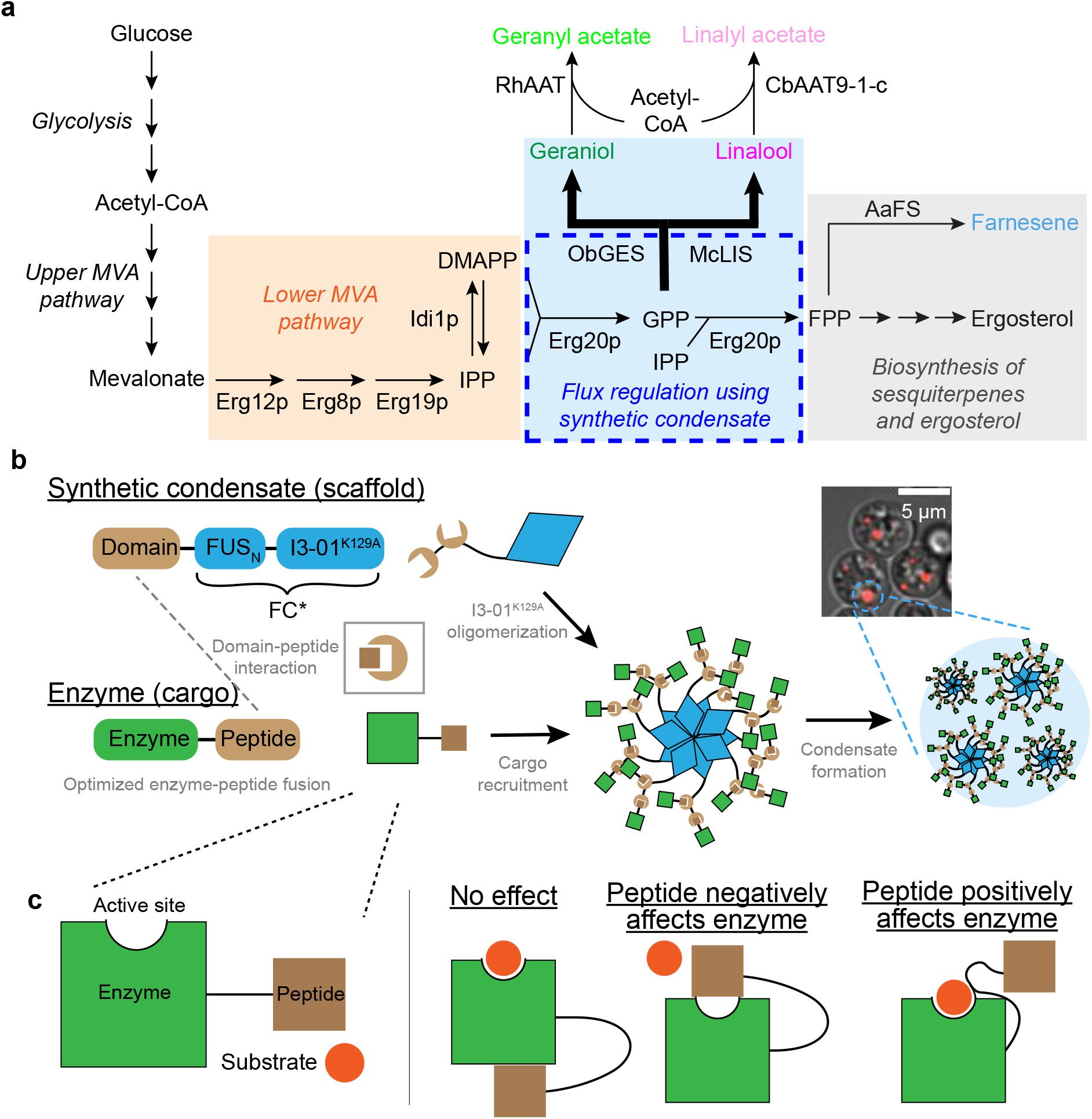
Overview of the monoterpene biosynthesis pathway in *Saccharomyces cerevisiae* and the application of synthetic condensate to regulate metabolic flux. **(a)** In the monoterpene biosynthesis pathway, the key precursor molecule mevalonate (MVA) is converted to GPP via the lower MVA pathway, composed of Erg12p, Erg8p, Erg19p, and Idi1p, and the bifunctional GPP/FPP synthase Erg20p. GPP is the common precursor molecule of monoterpenes, but Erg20p quickly converts GPP to FPP. In yeast, FPP is further converted to ergosterol, a molecule essential for yeast growth. Metabolite abbreviations: isopentenyl diphosphate (IPP), dimethylallyl diphosphate (DMAPP), geranyl diphosphate (GPP), farnesyl diphosphate (FPP). **(b)** Schematics of the synthetic condensate system, namely the Forever Corelet (FC^*^), and its functionalization with protein domain for the recruitment of peptide-fused enzymes. Representative image of yeast expressing FC* labeled with mCherry red fluorescent protein is shown. The scale bar represents 5 μm. **(c)** Flexible peptide tags required for recruitment to condensates may interact with the enzyme active site upon covalent fusion.

The presence of an endogenous MVA pathway makes *Saccharomyces cerevisiae* an attractive industrial host for monoterpene biosynthesis. Furthermore, *S. cerevisiae* has proven highly effective for the heterologous expression of cytochrome P450 enzymes, thereby enabling the biosynthesis of diverse monoterpene derivatives. These include monoterpene indole alkaloids with broad pharmaceutical and biological activities, such as the anti-cancer drug vinblastine^10,11^. However, GPP availability in *S. cerevisiae* remains low, as Erg20p predominantly functions as an FPP synthase rather than a GPP synthase^12^. In yeast, FPP is a key precursor for ergosterol biosynthesis, the principal sterol that governs plasma membrane permeability, fluidity, and protein function^13^. FPP synthase activity is essential for cell growth and maintaining sufficient intracellular levels of FPP is necessary for cell viability^14^. Heterologous expression of plant GPPS has not been shown to have a significant effect on monoterpene production.^15,16^ Due to the limited availability of GPP in yeast, reported monoterpene titers range between hundreds to thousands of mg/L^14,17^, whereas FPP-derived sesquiterpenes can reach tens to hundreds of g/L, enabling several successful industrial-scale applications^18–20^. To overcome this bottleneck, several mutations (F96W, N127W, and K197G) in the A1 site of Erg20p have been rationally engineered to slow the conversion of GPP to FPP, thereby enhancing monoterpene production^16,21^.

In addition to protein engineering strategies that enhance target molecule production, an emerging approach to further increase metabolic flux involves the colocalization of pathway enzymes into multienzyme complexes^22^. Natively, cells compartmentalize metabolic pathways within membraneless organelles such as purinosomes and G-bodies, which facilitate purine biosynthesis^23^ and glycolysis^24,25^, respectively. Inspired by such systems, we have previously developed a platform for building synthetic biomolecular condensates, termed Forever Corelets (FC)^26–28^, in which the N-terminal intrinsically disordered region (IDR) of human Fused in Sarcoma (FUSn) is fused to multimeric protein scaffolds, including the 24-mer human ferritin^29^ and the 60-mer synthetic scaffold protein, I3-01^K129A,30^. To functionalize FCs into synthetic membraneless organelles, we constructed a modular recruitment system by fusing protein-peptide interaction domains (PpIDs) to FUSn and the cognate PpID peptides to enzymes of interest^26^. In this work, we combine the triple effect of active site mutations in Erg20p, enhanced GPP synthase activity, and compartmentalization in synthetic membraneless organelles to improve the production of monoterpenes, including geraniol, linalool, and their esters by up to 5-fold, achieving geraniol titers exceeding 4 g/L. Using the PpID-FC system, we recruited different Erg20p variants and monoterpene synthases to synthetic organelles to channel the GPP intermediate toward monoterpene production. We serendipitously discovered that fusing short peptides, as short as two amino acids, to the C-terminus of Erg20p significantly boosts GPP synthase activity over FPP synthase activity by a previously unrecognized interaction between the C-terminus and active site of Erg20p. Determination of the steady-state kinetic parameters of Erg20p *in vitro* reveals that C-terminal fusions of short peptides lead to significant changes in catalytic efficiency, reversing the enzyme’s native activity from predominantly FPP synthase to GPP synthase. Overall, this study shows that synthetic organelles can significantly boost the productivity of compartmentalized metabolic pathways and, in particular, that short C-terminal peptide fusions in Erg20p enhance monoterpene production.

## Results

### C-terminal peptide fusions to yeast Erg20p improve monoterpene production

Previously, we characterized a suite of protein domain-peptide interaction pairs (PpIDs) to selectively enrich enzymes within synthetic organelles^26^. In this scheme, cognate peptides are directly fused to enzymes of interest, while the corresponding protein domains are fused to FC scaffolds (Fig 1b). To isolate the intrinsic effects of these flexible peptide tags on enzyme activity, we first examined their influence in the absence of compartmentalization within synthetic organelles (Fig 1c).

The first such tag we tested was the cognate PDZ peptide (PDZ_P_), which interacts with the PDZ domain (PDZ_D_) from α-syntrophin and mediates enzyme recruitment to PDZ-functionalized FCs^31^ (Fig 1b).^31^

As a control experiment, and prior to targeting these enzymes to synthetic organelles, we evaluated whether C-terminal fusions affect Erg20p activity by comparing geraniol production in strains expressing either the tagged or wild-type enzyme. To achieve this, *ERG20* in a previously developed mevalonate-overproducing *S. cerevisiae* strain (JCY51, which overexpresses both the upper and lower mevalonate pathways) ^32^ was replaced with *ERG20* variants carrying different C-terminal peptide fusions (Figure 2a). Additionally, to enable geraniol production from GPP, two copies of geraniol synthase from *Ocimum basilicum* (ObGES) were genomically integrated. To determine whether enhanced geraniol production correlates with reduced intracellular FPP levels, we quantified farnesene production as a proxy for intracellular FPP. To this end, a previously characterized mutant farnesene synthase from *Artemisia annua*, containing thirteen point mutations that confer an eight-fold increase in enzymatic activity relative to the wild-type enzyme (“AaFS_mut_”)^26,33^, was genomically integrated into all monoterpene-producing strains. The strain expressing Erg20p fused to a 43 amino acid-long tag containing the cognate PDZ peptide (PDZ_P_) and a Myc tag connected by two linker sequences (called Erg20p–Myc–PDZ_P_, strain yKX690, see Figure 2a and Supplementary Table 3) produced 172 ± 2 mg/L of geraniol, which is surprisingly 1.8-fold higher than the wild-type control strain (yKX607) expressing untagged Erg20p (Figure 2b). The same Erg20p–Myc–PDZ_P_ fusion strain (yKX690) also exhibits an 18% reduction in farnesene production compared to the control (yKX607), consistent with the C-terminal peptide fusion enhancing GPP synthase activity at the expense of FPP formation (Figure 2b).

**Figure 2.**
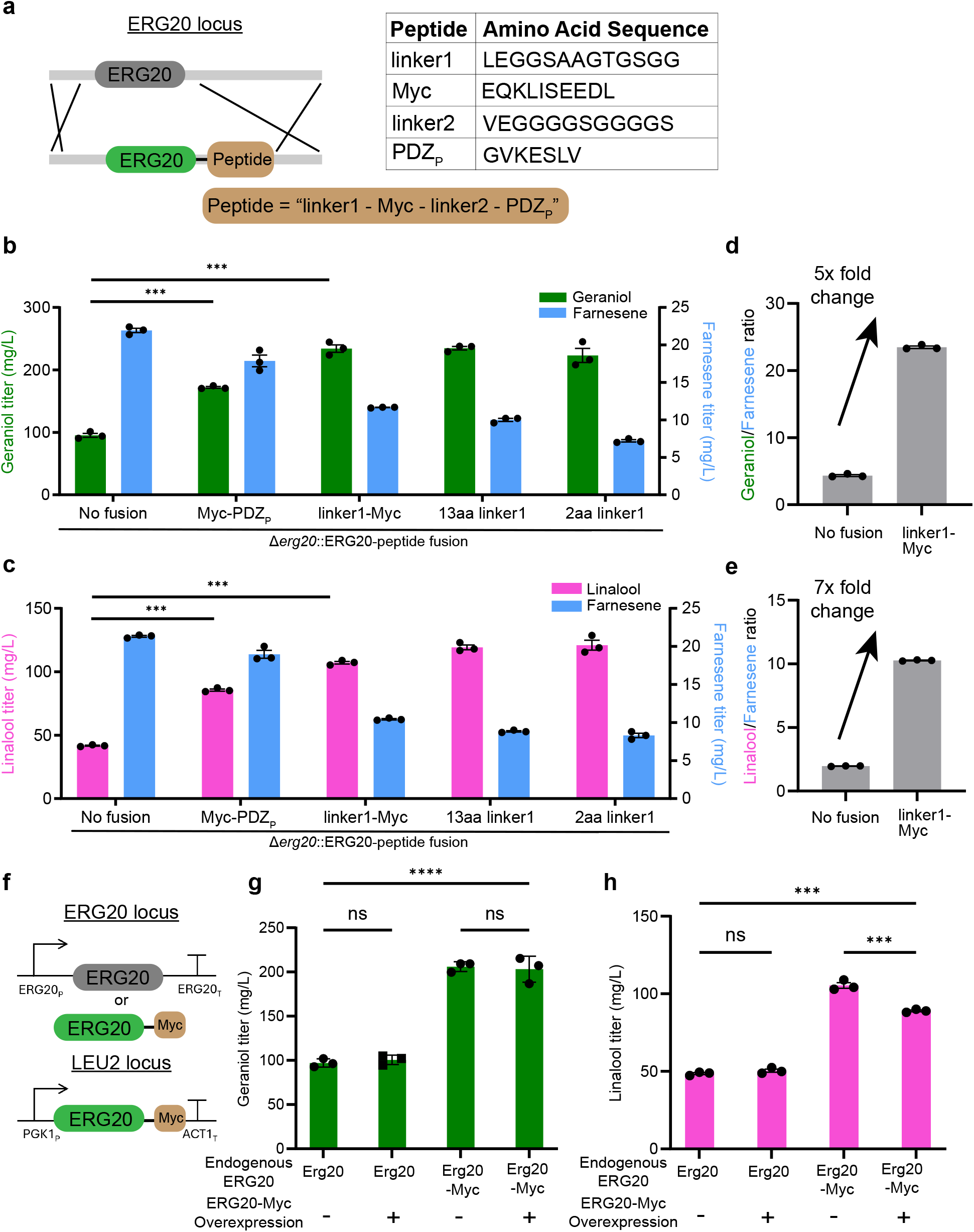
C-terminal fusion of short peptides to Erg20p improves monoterpene production. **(a)** Schematic representation of the CRISPR/Cas9-mediated homology recombination process to replace endogenous ERG20 gene with engineered ERG20–peptide fusions. The PDZ peptide (PDZ_P_) enables the enrichment of Erg20p in synthetic condensates functionalized with the PDZ domain. The corresponding peptide sequences are provided in the table. **(b)** Geraniol and farnesene productions in yeast strains where ERG20 has been replaced with ERG20–peptide fusions of varying lengths. **(c)** Linalool and farnesene production in yeast strains where ERG20 has been replaced by ERG20–peptide fusions of varying lengths.**(d)** Comparison of the geraniol-to-farnesene ratio in yeast expressing the Erg20p–linker1-Myc fusion from the endogenous *ERG20* locus compared to the wild-type control. **(e)** Comparison of the linalool-to-farnesene ratio in yeast expressing the Erg20p–linker1-Myc fusion from the endogenous *ERG20* locus compared to the wild-type control. **(f)** Schematic of the experimental design to determine whether ERG20– peptide fusions are dominant or recessive modifications. An additional copy of ERG20–Myc is expressed from the *LEU2* locus with or without the presence of endogenous Erg20p **(g-h)** Estimation of intracellular GPP levels in yeast overexpressing Erg20p–Myc, with and without endogenous Erg20p, using either geraniol (g) or linalool (h) as readouts. Error bars represent one standard deviation from three biological replicates. Statistics are calculated using a two-sided t-test. ns (non-significant); ^***^P < 0.001.

To ascertain whether this unexpected effect of the Erg20p–Myc–PDZ_P_ fusion reflects a general increase in intracellular GPP availability rather than a product-specific effect on geraniol synthesis, we tested its impact on the production of a second monoterpene, linalool. For this, we constructed linalool-producing strains by genomically integrating two copies of an engineered linalool synthase from *Mentha citrata* (t67McLIS^E343D, E352H^)^34^ into strains expressing either wild-type Erg20p or Erg20p–Myc–PDZ_P_, both also carrying AaFS_mut_ to report on FPP levels. Consistent with results from the geraniol-producing strains, the C-terminal Myc–PDZ_P_ fusion doubled linalool titers from 42 ± 1 mg/L to 86 ± 2 mg/L (strain yKX730) compared to the control expressing untagged Erg20p (strain yKX660) (Figure 2c). Similar to the geraniol producing strains, this increase in linalool production was accompanied by a proportional reduction in farnesene synthesis, confirming that the Myc-PDZ_P_ fusion redirects metabolic flux toward GPP formation at the expense of FPP.

Next, we sought to determine if the identity of the Erg20p tag was responsible for the observed increase in geraniol and linalool production. We replaced the PDZ_P_ peptide with a different cognate peptide, SH3_P_. The strain expressing Erg20p–Myc–SH3_P_ (strain yKX692) exhibited a comparable increase in geraniol titer relative to the untagged control (Figure S1). Fusion of a hemaglutinin (HA) peptide to the C-terminus of Erg20p also shifted preference towards GPP synthesis over FPP formation (Figure S2, strain yKX721), although the overall enzymatic activity was reduced as well. We concluded that enhancement in geraniol and linalool production was not specific to the Erg20p PDZ_P_ peptide tag. To explore the impact of the peptide length on this effect, we truncated the PpID entirely, leaving only the linker1 and the Myc tag. Surprisingly again, this much shorter construct further increased flux to geraniol over farnesene. To find the minimal tag length capable of causing this effect we continued to trim down the C-terminal peptide fusion. We made several truncations from the C-terminus of the tag, leaving behind peptides of 13 amino acids long, 7 amino acids long, and 2 amino acids long (Supplementary Table S1; respectively, yKX720,yKX793, yKX794). Surprisingly, all tested truncations of the linker1-Myc Erg20p fusion, including the addition of just two amino acids to the C-terminus, resulted in decreased FPP levels and enhanced geraniol and linalool production relative to the unfused control (Figure 2b, c). Notably, the optimal peptide fusion of 7 amino acids (sequence: LEGGSAA; strain yKX793) increased the geraniol titer by 2.9-fold to 278 ± 8 mg/L with a 10-fold improvement in the geraniol-to-farnesene ratio, and the linalool titer by 3.5-fold to 146 ± 5 mg/L (strain yKX798), followed by a 7-fold improvement in the linalool-to-farnesene ratio relative to the wild-type control strain (Figure 2b-e). Collectively, these results demonstrate that short C-terminal peptide fusions to yeast Erg20p can significantly improve monoterpene production by increasing GPP formation while reducing FPP accumulation.

Previously engineered Erg20p mutants have been shown to follow dominant negative inheritance patterns, and we sought to determine whether these C-terminal fusion Erg20p variants exhibit similar behavior. To determine whether replacing the endogenous *ERG20* with *ERG20*–peptide fusions is critical for the observed increase in monoterpene titers, we tested whether the fusion proteins act dominantly or recessively to wild-type Erg20p. We integrated an additional copy of the *ERG20*–Myc fusion into the *LEU2* locus of strains expressing either endogenous Erg20p or Erg20p–Myc from the native *ERG20* locus (Figure 2f) (strain yKX600, yKX601, yKX707, yKX778). The presence of residual endogenous Erg20p significantly reduced monoterpene titers, even in the presence of overexpressed Erg20p–Myc (Figure 2g, h), suggesting that the formation of Erg20p/Erg20-Myc heterodimers may underlie this inheritance pattern. In contrast, complete replacement of endogenous Erg20p with Erg20p–Myc was sufficient to enhance production by approximately 2-fold without further overexpression, indicating that Erg20p–Myc acts as a recessive, hypomorphic allele of Erg20p (Figure 2g, h). As the dominant wild-type Erg20p masks the DMAPP selectivity of the recessive Erg20p–Myc variant, removal of the endogenous ERG20 is necessary to redirect flux toward monoterpene synthesis. Together, these results demonstrate a generalizable strategy for fine-tuning pathway flux by introducing a recessive hypomorphic allele combined with deletion of the endogenous dominant gene, thereby circumventing both the lethality of complete gene deletion and the metabolic burden associated with overexpression.

### Only C-terminal peptide fusions enhance GPP synthase activity in Erg20p fusions

To determine whether the apparent increased intracellular GPP/FPP ratio observed in Erg20p peptide fusions is specific to the C-terminal fusions, we engineered Erg20p constructs with the same 13-residue linker1 sequence fused to either the N- or C-terminus of Erg20p (strain yKX720, yKX797). The N-terminal fusion did not significantly alter product distribution, whereas the C-terminal fusion significantly increased both geraniol and linalool titers at the expense of farnesene production (Figure S3a, b). These results demonstrate that C-terminal positioning of the peptide fusion is critical for shifting Erg20p selectivity toward GPP formation and promoting monoterpene synthesis. We then fused 6xHis tag to the N-terminus of Erg20p. The strain expressing 6xHis– Erg20p (yKX718) produced geraniol and farnesene titers comparable to the control strain expressing untagged Erg20p (Figure 3a), indicating that the N-terminal tag does not measurably affect enzyme function. In contrast, fusing a linker1–Myc sequence to the C-terminus of 6xHis– Erg20p increased geraniol titers 2-fold (strain yKX719), reaching 227 ± 13 mg/L, consistent with our previous findings (Figure 3a). These results show that the increased geraniol production due to the presence of the C-terminal fusion tag is preserved in the context of the N-terminal 6xHis tag.

**Figure 3.**
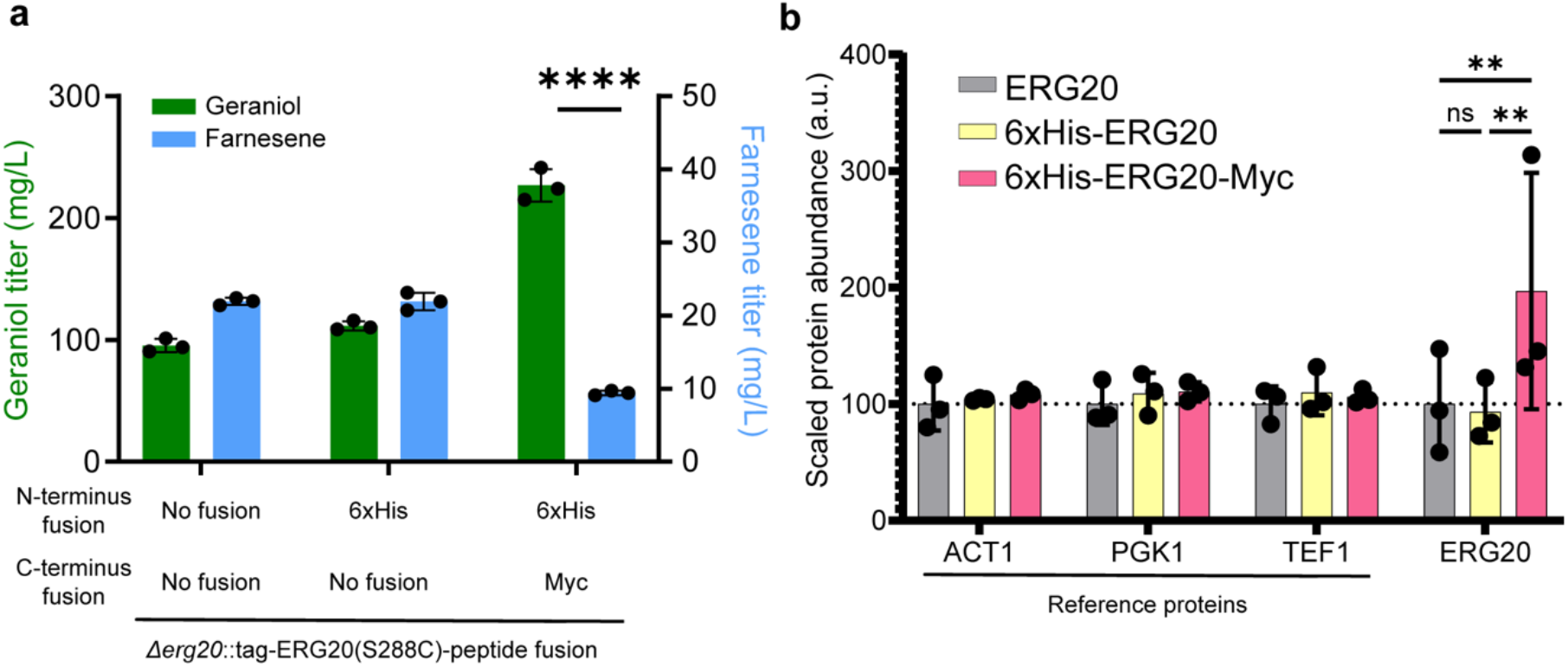
C-terminal fusion of short peptides improves intracellular accumulation of Erg20p. **(a)** Geraniol and farnesene production levels in yeast strains where the endogenous Erg20p is replaced with either N-terminally 6xHis tagged Erg20p or Erg20p–Myc. **(b)** Relative abundances of Erg20p and endogenous yeast reference proteins (encoded by ACT1, PGK1, and TEF1) as determined by whole-cell proteomics analysis. Abundance values were scaled such that the strain where wild-type *ERG20* is integrated has an average of 100. Error bars represent one standard deviation from three biological replicates. Statistics are calculated using a two-sided t-test. ns (non-significant); ^**^P < 0.01.

To assess how much of the observed effect of the C-terminal peptide fusion can be attributed to elevated Erg20p expression levels, we performed whole-cell proteomics to quantify intracellular 6xHis–Erg20p and 6xHis–Erg20p–Myc levels when expressed from the endogenous *ERG20* promoter and locus. We found that the abundance of 6xHis–Erg20p–Myc was higher than that of either 6xHis–Erg20p or untagged Erg20p controls (Figure 3b). Furthermore, we observed an overall upregulation of enzymes in both the upper and lower branches of the MVA pathway, as well as in the ergosterol biosynthesis pathway, in response to the C-terminal linker1–Myc fusion to Erg20p (Figure S4), consistent with reduced ergosterol biosynthesis^32,35^. This suggests that decreased intracellular FPP levels caused by the C-terminal fusion affect ergosterol biosynthesis.

### In vitro studies confirm that C-terminal peptide fusions improve GPP synthase of Erg20p

To dissect how C-terminal peptide fusions alter Erg20p activity independently of a cellular regulation, we performed steady-state kinetic analyses on purified enzymes. N-terminally His-tagged Erg20p variants were expressed in *E. coli*, purified by affinity chromatography (Figure S5), and analyzed to determine their kinetic parameters (Table 1; Figure S6). Consistent with our previous *in cellulo* observations, the C-terminal Myc fusion shifts the enzyme’s substrate preference from GPP to DMAPP (Table 1). The Myc-tag increased the apparent affinity (lower *K*_*M*_) for DMAPP by approximately twofold and decreased the affinity for GPP by roughly tenfold relative to the untagged enzyme. Together with modest increases in *k*_*cat*_ for both reactions, this resulted in an overall 3-fold increase in catalytic efficiency (*k*_*cat*_*/K*_*M*_) for DMAPP (Tukey HSD p-value = 0.002) and a 6.6-fold decrease for GPP (p-value = 0.02) compared with wild-type Erg20p (Table 1). This shift results in more than a 20-fold increase in the DMAPP/GPP preference ratio. The combined effects of accelerated GPP synthesis and attenuated GPP-to-FPP conversion account for the enhanced monoterpene production observed in strains expressing C-terminally tagged Erg20p.

**Table 1.** Steady-state kinetic parameters of Erg20p and Erg20p variants.

| | $K_M^{\text{DMAPP}}$ (mM) | $k_{\text{cat}}^{\text{DMAPP}}$ ($\text{s}^{-1}$) | $k_{\text{cat}}^{\text{DMAPP}}/K_M^{\text{DMAPP}}$ ($\text{mM}^{-1}\text{s}^{-1}$) |
| --- | --- | --- | --- |
| Erg20p | $0.21 \pm 0.05$ | $0.13 \pm 0.01$ | $0.59 \pm 0.15$ |
| Erg20p-Myc | $0.10 \pm 0.02$ | $0.19 \pm 0.01$ | $1.79 \pm 0.31$ |
| Erg20p(F96W) | $0.34 \pm 0.08$ | $0.10 \pm 0.01$ | $0.29 \pm 0.07$ |
| Erg20p(F96W)-Myc | $0.26 \pm 0.05$ | $0.11 \pm 0.01$ | $0.43 \pm 0.08$ |

| | $K_M^{\text{GPP}}$ (mM) | $k_{\text{cat}}^{\text{GPP}}$ ($\text{s}^{-1}$) | $k_{\text{cat}}^{\text{GPP}}/K_M^{\text{GPP}}$ ($\text{mM}^{-1}\text{s}^{-1}$) |
| --- | --- | --- | --- |
| Erg20p | $0.57 \pm 0.18$ | $3.53 \pm 0.37$ | $6.15 \pm 2.04$ |
| Erg20p-Myc | $5.89 \pm 2.63$ | $5.51 \pm 1.47$ | $0.93 \pm 0.49$ |
| Erg20p(F96W) | $12.82 \pm 6.45$ | $3.92 \pm 1.36$ | $0.31 \pm 0.19$ |
| Erg20p(F96W)-Myc | $3.54 \pm 0.54$ | $2.51 \pm 0.17$ | $0.71 \pm 0.12$ |

To assess how the C-terminal peptide fusion alters enzyme kinetics relative to the established GPP-favoring Erg20p mutant, F96W^16^, we quantified the FPP and GPP synthase activities of both Erg20p(F96W) and its fused variant, Erg20p(F96W)–Myc. Both variants were purified by affinity chromatography and assayed for steady-state kinetic parameters as previously described (Table 1; Figure S5 and S6). The presence of the F96W mutation appears to modulate the influence of the Myc tag, suggesting that the tag and mutation affect distinct structural regions of the active site. Consistent with earlier reports, the F96W mutation alone reduced catalytic efficiency (*k*_*cat*_*/K*_*M*_) for both substrates, primarily by slowing GPP-to-FPP conversion while modestly impairing DMAPP-to-GPP turnover.^16^ Compared with wild-type Erg20p, Erg20p(F96W) displayed significantly lower apparent affinity (higher *K*_*M*_) for both substrates (p-value _DMAPP_ = 0.02, p-value _GPP_ = 0.03). Adding the Myc tag to Erg20p(F96W) partially restored GPP synthase catalytic efficiency, such that there was no significant difference between the *k*_*cat*_*/K*_*M*_ of wild-type Erg20p and Erg20p(F96W)-Myc. The catalytic efficiency of Erg20p(F96W)–Myc for GPP as substrate remained similarly impaired, showing no significant difference from either wild-type Erg20p-Myc (p-value _GPP_ > 0.05) or Erg20p(F96W) (p-value _GPP_ > 0.05), indicating that the FPP synthase activity is comparably reduced across all variants. Overall, these results indicate that the C-terminal Myc fusion enhances GPP synthase efficiency in both wild-type and F96W backgrounds, suggesting that increased monoterpene production arises from a combined effect of improved GPP synthesis and reduced FPP formation.

### Structural basis for C-terminal tag effect on Erg20p GPP synthase activity

To elucidate the structural mechanism underlying the effect of C-terminal peptide fusion on Erg20p activity, particularly the significant reduction in GPP-to-FPP conversion, we employed a combined Alphafold2 and Rosetta modeling pipeline to investigate changes in the Erg20p active site. It has been previously shown that the F96W mutant sterically hinders the allylic site (A1), thereby driving preferential binding of the smaller DMAPP over the larger GPP (Figure S7a).^16^ The wild-type *S. cerevisiae* Erg20p model indicates that the C-terminus, which contains a conserved arginine residue, is located proximal to the enzyme’s active site (Figure 4b). Additionally, the model suggests that fusion of ‘linker1–Myc’ extends the C-terminal loop into the active site pocket proximal to the homoallylic IPP binding site (HA1), reducing its volume by 53% (Figure 4b and c). However, the addition of just two amino acids, leucine and glutamic acid (LE), to the C-terminus is sufficient to achieve a comparable reduction in HA1 site volume to that observed with the longer C-terminal fusion (Figure 4c). This finding is consistent with the equivalent enhancement in monoterpene production observed experimentally (Figure 2, strain yKX796). We hypothesize that changes in Erg20p activity correlate with a reduction in active site pocket volume, which hinders binding of the larger GPP molecule while still permitting binding of the smaller substrates DMAPP and IPP. Guided by this hypothesis, we used rational design to introduce targeted amino acid additions to the C-terminus, generating Erg20p variants spanning a range of predicted active site pocket volumes (Figure 4c). Notably, most designs extend the disordered C-terminus towards the HA1 site, thereby reducing the pocket volume, with the exception of the two leucine (LL) addition. Unlike all other fusions, which are predicted to lack defined secondary structure, the LL fusion is predicted to convert the loop region into a structured α-helix. This structural change results in minimal alteration of the active site pocket compared to wild-type Erg20p (Figure 4 c-e). As predicted by these models and consistent with our hypothesis, experimental expression of Erg20p variants with reduced active site volume results in a 2- to 3-fold increase in geraniol production, accompanied by a corresponding decrease in farnesene production (Figure 4f; left to right, strain yKX813, yKX821, yKX822, yKX810, yKX811). In contrast, the Erg20p-LL (strain yKX820) variant produced 53% less geraniol compared to the wild-type (Figure 4d), suggesting that the propensity of these additional residues to form an α-helix at the C-terminus prevents the reduction in HA1 volume, and that the coiled conformation of the C-terminus in the wild-type is important for enzymatic activity. Taken together, our findings indicate that the coiled C-terminus of Erg20p influences the active site pocket volume through steric hinderance at the HA1 site. Thus, by engineering amino acid fusions to the C-terminus, we can modulate the extent of intrusion at the HA1 site, thereby affecting the active site pocket volume and providing a means to fine-tune substrate binding.

**Figure 4.**
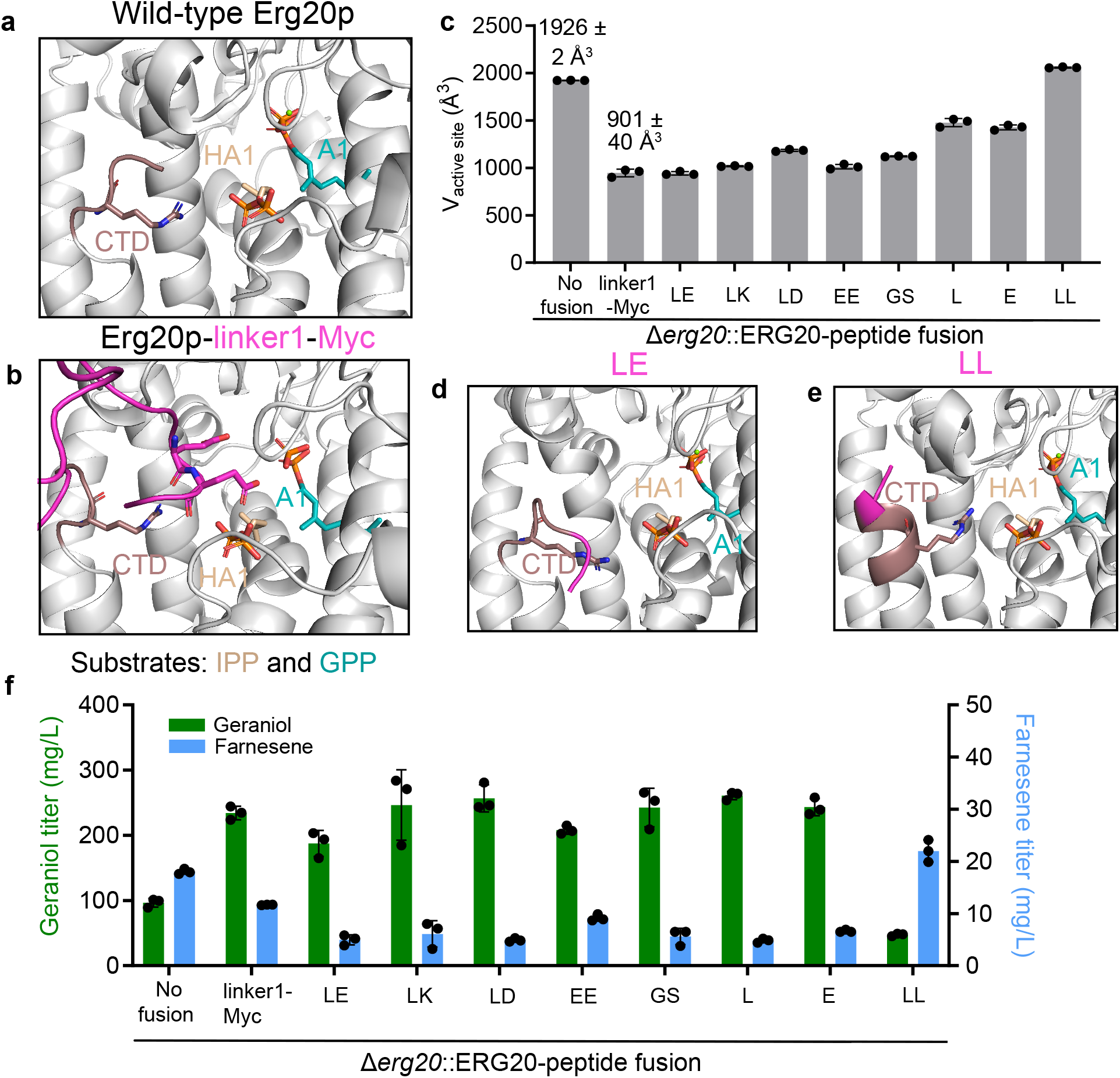
Impact of C-terminal fusion on Erg20p active site pocket volume and geraniol production. **(a)** Structural model of wild-type Erg20p and **(b)** the Erg20p-linker1-Myc fusion, showing the disordered C-terminus of *S. cerevisiae* Erg20p (brown) positioned near the active site. Fusion of the “linker1-Myc” peptide (magenta) to the C-terminus reduces the active site pocket volume. The arginine side chain at the C-terminus of wild-type Erg20p and all amino acid side chains within the active site of the linker1-Myc fusion are displayed. Bound substrates IPP (cream) and GPP (cyan) occupy the homoallylic HA1 site and allylic A1 site, respectively. **(c)** Rational design of C-terminal amino acid fusions to Erg20p to generate a range of active site pocket volumes. **(d)** Structural model of the rationally designed Erg20p-LE variant and **(e)** Erg20p-LL variant, showing how amino acid fusions alter the coil propensity of the C-terminus. The disordered C-terminus of wild-type *S. cerevisiae* Erg20p is shown in brown, and the fused amino acids are shown in magenta. **(f)** Geraniol and farnesene production in yeast strains in which native Erg20p has been replaced with rationally designed Erg20p fusion variants.

However, the cross-effects arising from simultaneous mutation of the A1 site and C-terminal peptide fusion are complex. While the F96W mutation decreases the active site volume to a similar extent as the Myc tag when introduced individually (Figure 4c, S7b), in the presence of both F96W and the Myc tag, our models suggest the Myc tag flexibly rearranges the HA1 site such that E373 coordinates the pyrophosphate groups of IPP, resulting in an overall increase in active site volume (Figure 4h). Thus, Erg20p(F96W)–Myc recovers much of the pocket volume of unfused wild-type Erg20p (Figure 4c, S7a and S7b), potentially contributing to the decreased apparent affinity (higher *K*_*M*_) for DMAPP relative to Erg20p–Myc, as well as the recovery of apparent GPP affinity in Erg20p(F96W)–Myc compared with either Erg20p(F96W) or Erg20p–Myc (Table 1). Together, these findings point to a structural basis underlying the production trends observed in yeast and the activity measured for purified Erg20p.

### Employing the Erg20p–Myc strain with improved GPP production as a chassis for monoterpene biosynthesis

Given the significant improvement in GPP synthase activity conferred by Erg20p–Myc, we tested whether a strain expressing Erg20p–Myc could serve as a chassis for improved monoterpene biosynthesis. For geraniol production, increasing the gene copy number of geraniol synthase (ObGES) from 2 to 6 did not result in a further increase in geraniol titers (Figure 5a; strain KX806, yKX807, yKX760), indicating that ObGES expression is not limiting under these conditions. In contrast, overexpression of the lower mevalonate pathway in the strain expressing 6 copies of ObGES led to a 67% increase in geraniol production, from 323 ± 9 mg/L to 538 ± 9 mg/L (Figure 5a, strain yKX642). To further derivatize geraniol, we integrated a single copy of the *Rosa hybrida* alcohol acyltransferase (RhAAT). The resulting strain (yKX643) produced 1040 ± 30 mg/L of geranyl acetate, representing the highest titer reported to date in *S. cerevisiae*, with nearly complete conversion of geraniol (Figure 5a).

**Figure 5.**
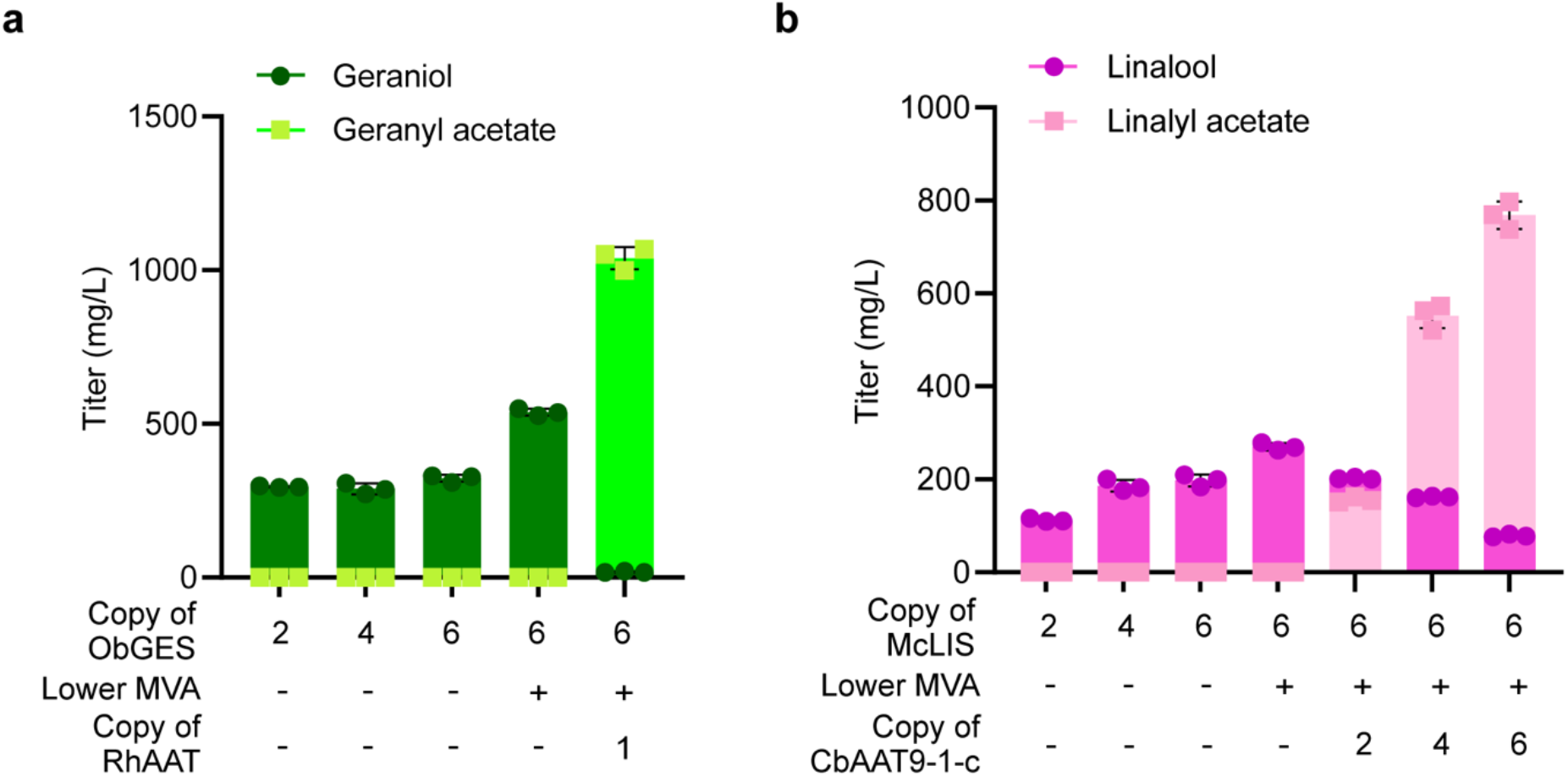
Production of diverse monoterpene molecules in yeast expressing endogenous Erg20p-Myc. **(a)** Optimization of geraniol production by overexpressing additional copies of ObGES and the lower mevalonate pathway. Geraniol can be further converted to geranyl acetate by RhAAT. **(b)** Optimization of linalool production by overexpressing additional copies of McLIS^E343D, E352H^ and the lower mevalonate pathway. Linalool can be further converted to linalyl acetate by CbAAT9-1-c. Error bars represent one standard deviation from three biological replicates.

Similarly, integrating additional copies of McLIS^E343D, E352H^ (6 copies in total) and overexpressing the lower mevalonate pathway (strain yKX734) increased linalool production to 270 ± 6 mg/L, to our best knowledge, the highest titer reported in *S. cerevisiae* shake flask cultures (Figure 5b)^36^. Overexpression of *Citrus bergamia* alcohol acyltransferase (CbAAT9-1-c)^37^ further enabled conversion of linalool into linalyl acetate. A strain carrying 6 genomic copies of CbAAT9-1-c (yKX737) produced 768 ± 24 mg/L of linalyl acetate with 79 ± 2 m^37^g/L of residual linalool. Surprisingly, our best linalyl acetate-producing strain yielded a molar equivalent of 4.4 mM linalool in total, a significant 2.5-fold increase over the 1.8 mM linalool produced by the strain without CbAAT9-1-c overexpression (Figure 5b). This finding is consistent with previous observations for geranyl acetate and suggests that esterification of monoterpenes could be a generalizable strategy to improve monoterpene production by alleviating product toxicity^38–40^. In conclusion, we have demonstrated that the Erg20p–peptide fusion enzyme supports high-level production of four different monoterpenes (i.e., geraniol, linalool, geranyl acetate, and linalyl acetate) and shown that yeast expressing Erg20p–Myc can be used as a chassis for monoterpene biosynthesis.

### Regulation of monoterpene flux using synthetic organelles

We next examined whether monoterpene production could be further enhanced by assembling pathway enzymes within engineered biomolecular condensates. We hypothesized that colocalizing Erg20p with geraniol synthase inside synthetic organelles would increase the likelihood of efficient channeling of GPP toward geraniol formation, thereby reducing its conversion to FPP or diffusion out of the condensate. Prior theoretical and experimental studies have shown that clustering enzymes at metabolic branch points can effectively direct shared intermediates toward a desired pathway, thereby enhancing product yields (Figure S8)^41^. However, the same theoretical framework predicts that colocalizing enzymes in a linear pathway (e.g., Erg20p catalyzing two consecutive reactions) can also accelerate intermediate turnover to FPP, provided that the catalytic rate exceeds the rate of intermediate diffusion away from the enzyme cluster.

We first validated that colocalization affects enzyme competition for the shared GPP intermediate. To determine if the FPP synthase activity of Erg20p–Myc–PDZ_P_ competes with ObGES–PDZ_P_ within condensates, we examined whether localizing Erg20p–Myc–PDZ_P_ alone to synthetic condensates using PDZ domain-functionalized “Forever Corelets” (FC) was sufficient to increase intracellular FPP levels (Figure 6a, left panel). An increase in farnesene production indicates accelerated GPP-to-FPP conversion. Indeed, localization of Erg20p–Myc–PDZ_P_ within synthetic condensates led to a 58% increase in farnesene titer compared to the nonlocalized control (1.09 ±0.06 vs 0.67 ± 0.05 mg/L/OD_600_) (Figure 6b, strain yKX638 and yKX639), confirming that Erg20p– Myc–PDZ_P_ would compete with ObGES–PDZ_P_ for GPP within the condensate. To leverage synthetic organelles as a platform for increasing flux toward geraniol, it is therefore necessary to reduce the FPP synthase activity of Erg20p–Myc–PDZ_P_.

**Figure 6.**
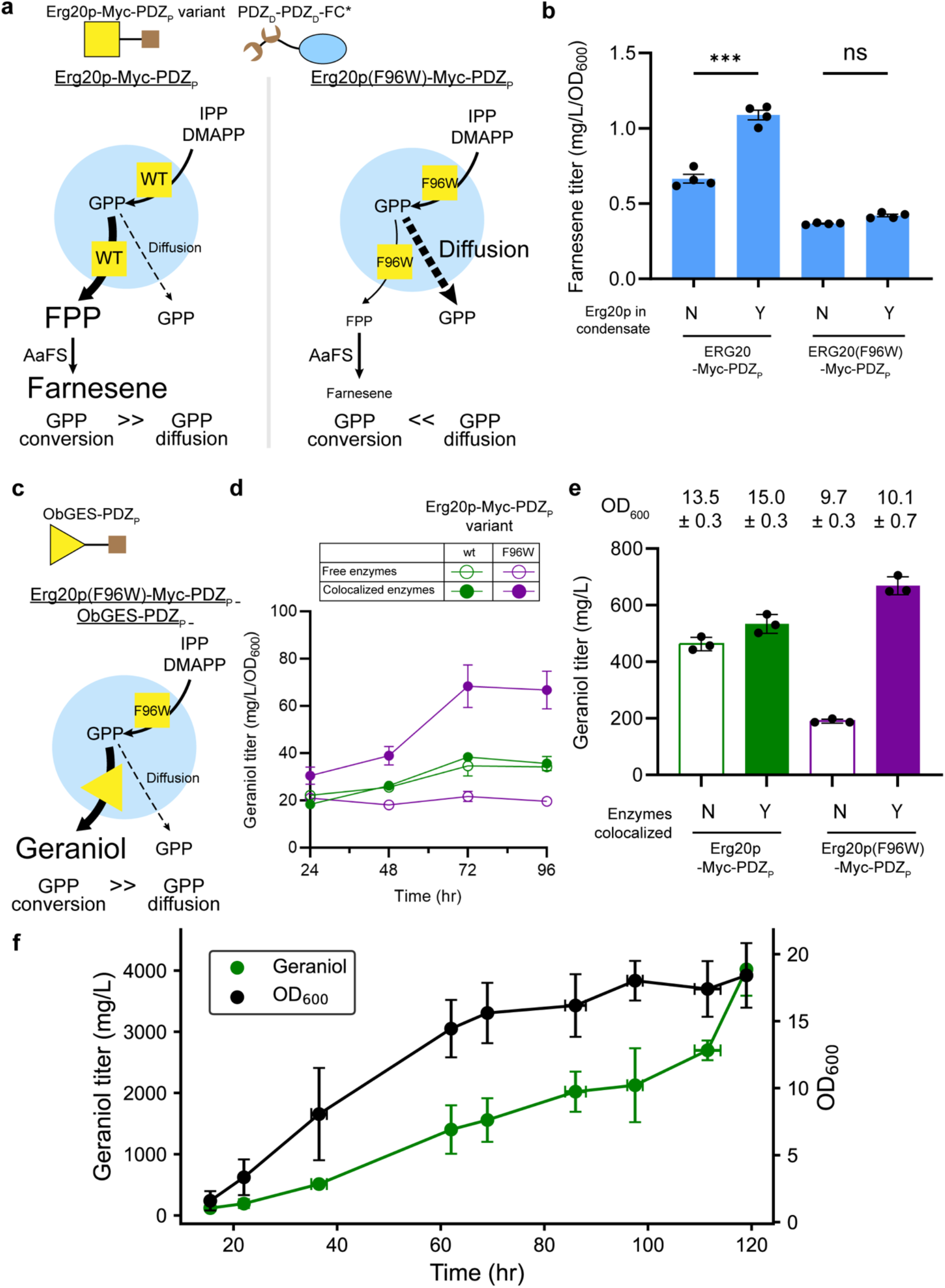
Enhancing geraniol production via synthetic condensate-mediated colocalization of Erg20p and geraniol synthase. **(a)** Schematic depicting the experiment designed to test the hypothesis that Erg20p–Myc–PDZ_P_ can compete with ObGES–PDZ_P_ for the shared GPP intermediate within Forever Corelet (FC*) condensates. Erg20p–Myc–PDZ_P_, with or without the F96W mutation, is localized within the condensates, and the resulting change in intracellular FPP levels is measured using farnesene as a readout. The F96W substitution reduces the FPP synthase activity of Erg20p and slows down GPP-to-FPP conversion to allow more GPP to diffuse away from the condensates. **(b)** OD_600_-nornalized farnesene titer in yeast strains expressing Erg20p–Myc–PDZ_P_, either localized within condensates or diffused, with and without the F96W mutation (n=3 biological replicates). **(c)** Schematic illustrating the colocalization of Erg20p and ObGES within Forever Corelet (FC*) synthetic condensates using protein domain-peptide interaction pairs to streamline geraniol production. **(d)** Time-course measurements of OD_600_-normalized geraniol production, comparing strains with and without colocalized Erg20p–Myc–PDZ_P_ and ObGES– PDZ_P_, as well as with or without the F96W mutation in Erg20p (n=3 biological replicates). **(e)** Final OD_600_ values and geraniol titers of yeast strains from panel (b) after 96hr of fermentation. **(f)** Production of geraniol and cell density in a 2L fed-batch bioreactor (n=2 biological replicates). Error bars represent one standard deviation. Statistics are calculated using a two-sided t-test. ns (non-significant); ^*^P < 0.05; ^***^P < 0.001.

To reduce flux from GPP to FPP within condensates, we introduced Erg20p(F96W)–Myc–PDZ_P_. We had previously shown *in vitro* that Erg20p(F96W)–Myc–PDZ_P_ exhibits lower catalytic efficiency for GPP consumption than Erg20p–Myc (Table 1), leading us to expect reduced flux toward FPP within condensates (Figure 6a, right panel). We found that a single copy of Erg20p(F96W)–Myc–PDZ_P_ expressed from the endogenous *ERG20* locus is sufficient to support yeast growth, although final OD_600_ was reduced by 28% compared to strains expressing Erg20p– Myc–PDZ_P_. As expected, Erg20p(F96W)–Myc–PDZ_P_ also decreased FPP accumulation (0.37 ±0.005 vs 0.67 ± 0.05 mg/L/OD_600_) (Figure 6b, strains yKX640 and yKX641), consistent with a slower rate of GPP to FPP conversion in the mutant than the rate of GPP diffusion out of the condensate (Supplementary Note 1).

Consistent with the observed differences in FPP flux, colocalization of Erg20p variants with ObGES–PDZ_P_ in condensates revealed contrasting outcomes (Figure 6c). Colocalization of Erg20p(F96W)–Myc–PDZ_P_ with ObGES–PDZ_P_ increased geraniol titers 3.5-fold compared to the corresponding free enzyme control strains (Figure 6d, e, strains yKX712 and yKX713). In contrast,colocalization of wild-type Erg20p–Myc–PDZ_P_ with ObGES–PDZ_P_ did not significantly affect geraniol production (Figure 6d, e, strains yKX710 and yKX711). We attribute this difference to the higher FPP synthase activity of wild-type Erg20p-Myc, which limits the benefit of colocalization by continuing to channel GPP toward FPP. By comparison, Erg20p(F96W)-Myc has reduced FPP synthase activity (Table 1), allowing colocalization with ObGES to more effectively divert GPP flux toward geraniol and thereby increase geraniol titers.

We observed similarly divergent trends in geraniol production under free enzyme conditions for wild-type Erg20p–Myc–PDZ_P_ and Erg20p(F96W)–Myc–PDZ_P_ (strains yKX711 and yKX713). In the absence of condensate recruitment, Erg20p(F96W)–Myc–PDZ_P_ resulted in a 58% decrease in geraniol titers compared to the corresponding dispersed wild-type Erg20p–Myc–PDZ_P_. This result is consistent with Erg20p(F96W)–Myc exhibiting a four-fold lower catalytic efficiency for DMAPP than Erg20p–Myc (Table 1), suggesting that reduced DMAPP conversion limits GPP availability under these conditions. While previous studies have reported increased geraniol production in yeast strains expressing free Erg20p(F96W) relative to wild-type Erg20p^16^, we attribute this discrepancy to the cross-effects of the codon 96 mutation and the C-terminal Myc tag, which in our system doubles the *k*_*cat*_*/K*_*M*_ with respect to GPP (Table 1).

Both in the recruited and dispersed conditions (strains yKX712 and yKX713, respectively), growth of the Erg20p(F96W)–Myc–PDZ_P_ strain was slower than that of the corresponding wild-type Erg20p-Myc–PDZ_P_ strain, consistent with previous reports of the Erg20p(F96W) mutant^42^. Reduced intracellular FPP levels (Figure 6b) may impair flux through the downstream ergosterol pathway, thereby affecting cell growth. In addition, accumulation of upstream intermediates such as IPP and DMAPP can exert cytotoxic effects^43,44^ and inhibit other enzymes in the mevalonate pathway^45^. Notably, excess FPP has also been reported to inhibit Erg20p activity,^42^ highlighting the importance of balanced intermediate levels. In strains where Erg20p(F96W)–Myc–PDZ_P_ and ObGES–PDZ_P_ are colocalized, OD-normalized geraniol titers continued to increase between 24 h and 72 h of fermentation, indicating that flux toward geraniol production is enhanced through proximity channeling of GPP within condensates (Figure 6d).

Having observed a marked increase in geraniol titers in shake-flask cultures, we next sought to scale production in a 2-L fed-batch bioreactor. We employed a prototrophic version of strain yKX712 (yKX786), in which Erg20p(F96W)–Myc–PDZ_P_ and ObGES–PDZ_P_ are colocalized within condensates. Cells were grown in SC medium supplemented with glucose. During fed-batch fermentation, both geraniol and biomass accumulated steadily, reaching a final geraniol titer of 4.04 ± 0.54 g/L and a final OD_600_ of 17.70 ± 1.78 after 120 hours (Figure 6f). These titers are comparable to the highest geraniol levels reported in yeast (5.52 g/L)^46^, but are achieved with significantly enhanced productivity: 0.0340 ± 0.0045 g/L/h reported here versus 0.0072 g/L/h.

In addition, we demonstrated that in geranyl acetate-producing yeast, colocalization of Erg20p– Myc–PDZ_P_, with or without the F96W substitution, with ObGES–PDZ_P_, improves geranyl acetate titers (Figure S10). This finding highlights the broader applicability of synthetic organelles for enhancing the production of monoterpene-derived compounds. Together, these results demonstrate that synthetic organelles complement protein engineering strategies by enabling spatial control over metabolic flux. While reducing FPP synthase activity alone introduces upstream bottlenecks, colocalization of Erg20p variants with downstream enzymes at the GPP branch point redirects flux toward desired products while mitigating the accumulation of toxic intermediates.

## Discussion

Monoterpene biosynthesis in microbial hosts offers a sustainable route to industrially valuable compounds; however, product titers have been historically constrained by competition for shared metabolic precursors. In this study, we address these limitations by integrating enzyme engineering with pathway compartmentalization within synthetic organelles, thereby redirecting metabolic flux from sesquiterpene toward monoterpene production.

Regarding enzyme engineering, we serendipitously found that fusing short peptides to the C-terminus of yeast Erg20p markedly alters its catalytic behavior – an effect that can be recapitulated by the addition of even a single amino-acid, provided it promotes a flexible C-terminus. Short peptide tags are generally considered to have minimal impact on enzyme activity^47^, yet growing evidence shows that they can modulate protein behavior in context-dependent manners. Peptide fusions have been reported to improve proteins folding, solubility, and stability^48,49^; as well as impart novel catalytic activity^48–50^, and increase the abundance or activity of rate-limiting metabolic enzymes *in vivo*^51,52^. Consistent with previous observations that GPP accumulates only at trace levels in yeast^16^, we found that wild-type Erg20p exhibits a 10-fold higher catalytic efficiency (*k*_*cat*_/*K*_*M*_) for GPP consumption than for GPP formation (Table 1). In contrast, C-terminal peptide fusions (e.g., linker-Myc) to Erg20p invert this substrate preference, yielding a 2-fold higher *k*_*cat*_/*K*_*M*_ for GPP formation relative to GPP consumption. According to molecular models, this kinetic shift arises from induced changes in the active site architecture (Figure 4) and correlates with the substantial increase in monoterpene titers observed in strains expressing Erg20p-peptide fusions (Figure 2).

Colocalization of metabolic enzymes is a well-established strategy for enhancing pathway efficiency by minimizing the loss of intermediates to competing reactions^53–56^. In plants, enzymes involved in monoterpene biosynthesis are naturally colocalized within membrane-bound plastids, effectively sequestering the GPP pool from cytosolic sesquiterpene pathways^57^. Beyond membrane-bound compartments, membraneless multi-enzymatic assemblies, or metabolons, have been proposed to dynamically organize metabolic pathways in nature and facilitate substrate channeling, thereby enhancing the biosynthesis of isoprenoids^58^ and other plant natural products^59^. Inspired by these natural architectures, we adopted our modular synthetic condensate platform^26^ to colocalize engineered Erg20p and geraniol synthase (ObGES) with the goal of further enhancing geraniol production. Reaction-diffusion models predict that the relative reaction rates of competing enzymes determine the extent of intermediate channeling within a metabolon, with maximal product formation achieved when the competing reaction is slowed^41^. Consistent with this model, strains with condensates where ObGES and Erg20p(F96W)-Myc are colocalized have significantly enhanced geraniol production relative to strains where ObGES and Erg20p-Myc are colocalized in condensates. Erg20p(F96W)-Myc exhibits a substantially reduced DMAPP and GPP turnover (Table 1) and so colocalization in condensates with ObGES accelerates GPP conversion (Figure 6b). Together, these results demonstrate a synergistic relationship between enzyme engineering and spatial organization, demonstrating that post-translational control of enzyme localization can be combined with kinetic tuning to optimize pathway flux under a reaction diffusion regime.

With these findings, this study establishes the use of synthetic membraneless organelles derived from protein phase separation as a feasible strategy in metabolic engineering to achieve meaningful titers of products of interest. Many challenges in metabolic engineering, including intermediate toxicity and metabolite loss to competing reactions, can be mitigated through enzyme colocalization. While previous studies have demonstrated proof-of-concept applications of synthetic organelles to metabolic engineering, thus far they have only achieved relatively modest improvements in titers^53,60,61^. Here we show that a strain co-localizing geranyl synthase and an engineered GPP synthase with carefully tuned kinetics within synthetic organelles achieves titers comparable to the highest ever reported in yeast, with a four-fold improvement in productivity (30 mg/L-h vs 8 mg/L-h)^46^. Drawing on lessons from this and prior studies, several limitations remain in the applicability of synthetic membraneless organelles for metabolic engineering.

Because they lack membranes to retain metabolites and isolate them from competing reactions in the cytosol, synthetic organelles formed by protein condensates can enhance targeted metabolic pathways only under certain kinetic constraints. Specifically, downstream targeted enzymatic reactions must outcompete both competing pathways and substrate diffusion within the condensate, ensuring that intermediates are converted before escaping the organelle^41^. The first criterion was manifest when colocalizing ObGES with Erg20p-Myc, which has continued competition between GPP and FPP synthesis (Figure 6b), resulting in only modest improvements in geraniol synthesis. In this regard, our results are even more remarkable in the sense that the competing reaction is also compartmentalized in the same condensate (inevitable by the nature of this Erg20, which has both geranyl synthase and farnesyl synthase activity). In most other pathways, however, competing reactions reside outside the synthetic organelle^26,53,62^, and therefore the effect of compartmentalization would be expected to be even more pronounced. In addition to slowing down or spatially separating competing reactions, the activity of the downstream enzymes may also be increased by the local environment of condensates^63–65^. Furthermore, the diffusivity of intermediates^66^, metabolites^67^ or cofactors^68^ may also be reduced by designing condensates with increased apparent viscosities to these small molecules. These additional features would be expected to further increase geraniol production by these monoterpene-biosynthetic synthetic organelles.

In conclusion, this study shows that synthetic organelles driven by protein phase separation can be effectively harnessed for metabolic engineering to achieve competitive production levels. When combined with enzyme engineering to optimize kinetics, the use of synthetic organelles to spatially segregate metabolic reactions is a compelling platform for controlling metabolic flux. Beyond enhancing the biosynthesis of monoterpenes and higher-value monoterpene-derived products, such as monoterpene indole alkaloids, in yeast, this strategy may be applied to diverse metabolic pathways in yeast and potentially other microbial hosts.

## Materials and Methods

### Plasmid Construction

All integration (*LEU2, HIS3, TRP1*) and 2μ plasmids were created based on the pJLA vectors (Supplementary Table S2), using either the restriction digest and ligation method with T4 DNA ligase (NEB) or the In-Fusion HD cloning kit (Takara Bio). Additionally, the EasyClone2.0 vector pCFB2225 (a gift from Irina Borodina, Addgene plasmid # 67553) was used to integrate gene constructs into the XII-2 locus^69^. The following restriction enzymes were used for cloning: MreI (Thermo Fisher Scientific), BspEI (NEB), SpeI-HF(NEB), BamHI-HF(NEB), NheI-HF (NEB), NotI-HF(NEB), AgeI-HF(NEB), AscI (NEB), and XhoI (NEB).

Promoters (CCW12p and HHF2p) and terminators (TDH1t, PGK1t, ENO2t, and ENO1t) that are not in the pJLA vectors were obtained from a published yeast toolkit on Addgene (Kit #1000000061), a gift from John Dueber^70^. *ERG20* in all plasmids refers to S288C *ERG20* variant. The C-terminal peptide modifications are introduced via oligo annealing followed by In-Fusion cloning. All oligos and primers are purchased from Integrated DNA Technologies (IDT). Recombinant DNA of heterologous enzymes (AaFS_mut_, RhAAT, McLIS, and CbAAT9-1-c) were codon-optimized for *S. cerevisiae* using IDT codon optimization tool and purchased as gBlocks® gene fragments from IDT. The AaFSmut sequence was based on a previously reported *Artemisia annua* variant ‘FS_C_8’, with mutations F11S, V24D, L18I, M35T, T196S, Y288F, T319S, R348K, T385A, I434T, T446N, I460V, V467I^33^. The synthetic condensate (domain fused FC*) constructs are obtained from previous studies^26,28^.

The cloned plasmids were transformed into Stellar *E. coli* competent cells (Takara Bio), plated on Luria-Bertani (LB) plates containing 100 μg/ml ampicillin, and incubated at 30°C overnight. Single colonies carrying the correct plasmid constructs were identified by colony PCR using OneTaq® Hot Start Quick-Load® 2X Master Mix (NEB), and plasmids were purified from overnight culture in LB containing 150 μg/ml ampicillin at 37°C. All cloned plasmids were verified by Sanger sequencing (Genewiz) or Whole Plasmid sequencing (Plasmidsaurus).

### Yeast Transformation and Fermentation

Integration plasmids were linearized with PmeI (NEB) or SwaI (NEB) and transformed into yeast using the standard lithium acetate method^71^. *S. cerevisiae* CEN.PK2-1C (MATa, ura3-52, trp1-289, leu2-3112, his3Δ1, MAL2-8c, SUC2) is the origin for all the engineered strains in this study (Supplementary Table S3).

Transformants were selected on synthetic complete (SC) agar plates containing 2% (w/v) glucose and lacking the appropriate amino acid for auxotrophic marker selection. To verify integration of the linearized DNA construct into the yeast genome, single colonies were isolated and screened using Phire Tissue Direct PCR Master Mix (Thermo Fisher Scientific). Positive colonies were subsequently evaluated in high-cell-density fermentations using 24-well plates. Yeast cultures were grown overnight in 1 mL of SC medium containing 2% (w/v) glucose and lacking the appropriate amino acid for selection. After 24 h of incubation at 30°C with shaking at 200 rpm, plates were centrifuged at 1000 rpm for 5 min. The supernatant was discarded, and cell pellets were resuspended in 1 mL of SC medium containing 2% (w/v) glucose, followed by the addition of a 20% (v/v) dodecane overlay (200 μL, Sigma Aldrich). Cultures were then incubated at 30°C for 48 hours with shaking at 200 rpm. At the end of fermentation, monoterpene production was quantified as described below. The highest producing colonies were stored as 20% (v/v) glycerol stocks at −80°C.

To compare strain performance, yeast strains were streaked from glycerol stocks onto yeast extract peptone dextrose (YPD) agar plates and incubated at 30°C until single colonies were visible. Individual colonies were then used to inoculate 1 mL of SC-HIS-LEU-TRP dropout medium containing 2% (w/v) glucose and a 20% (v/v) dodecane overlay (200 μL) in glass tubes, followed by overnight growth. The following day, overnight cultures were used to inoculate 3 mL of SC medium containing 2% (w/v) glucose and a 20% (v/v) dodecane overlay (600 μL) in 14 mL Falcon test tubes to a starting OD_600_ of 0.2. Cultures were then incubated at 30°C for 96 h with shaking at 265 rpm.

The wild-type, “no linker” (Figures 2-5) condition corresponds to the S288C *ERG20* variant integrated in strain JCY51 (CEN.PK2-1C background), replacing the endogenous CEN.PK2-1C *ERG20*. “Endogenous *ERG20*” (Figures 2-5) refers to the S288C *ERG20* integrated at the native *ERG20* locus, whereas “overexpressed *ERG20*” (Figures 2) denotes the S288C *ERG20* integrated at the *LEU2* locus. Where indicated, ObGES or 67McLIS^E343D, E352H^ were integrated at the *LEU2* locus; AaFS_mut_, was integrated at the *HIS3* locus.

### CRISPR/Cas9 Assisted Gene Editing

All monoterpene-producing strains with modified endogenous ERG20 in our study are derived from a mevalonate over-producing strain (JCY51)^32^. The Ellis lab CRISPR/Cas9 toolkit^72^ (a gift from Tom Ellis, Addgene) was used to endogenously tag the *ERG20* gene. The sgRNA sequences were designed to target the CEN.PK2-1C *ERG20* coding region (5’-ATGTTCTTGGGTAATCAACA<u>AGG</u>). The upstream and downstream regions of the target gene of interest were amplified from the yeast genome to construct the repair template. Colonies were screened and sequence verified as described above. To cure the Cas9 plasmid, colonies were cultured overnight in YPD media with 2% (w/v) glucose and then plated onto a YPD agar plate. To cure Cas9 plasmid with URA3 marker, overnight culture was plated onto YPD plate supplemented with 1g/L 5-fluoroorotic acid.

### Whole-cell Proteomics

Starter yeast cultures were grown overnight in 3 mL of YPD medium at 30°C and used to inoculate 1 L of YPD to an initial OD_600_ of 0.2. Cultures were incubated at 30°C for 48 hours with shaking at 220 rpm. Cells were harvested by centrifugation at 7500 x g for 20 minutes at 4°C, and the supernatant was discarded. Pellets were resuspended in 10 mL of phosphate buffered saline (PBS), flash-frozen in dots in liquid nitrogen, and subsequently disrupted at cryogenic temperatures using a Cryo-Mill (Spex sample prep^®^) system with 14 cycles of 2 min grinding (with impact frequency set at 13 cycles per second) followed by 3 min cooling. Lysed cells were stored in 50 mL conical tubes at −80°C.

Before trypsin digestion, the total protein content of samples was determined using the Bradford assay by measuring the absorbance of Coomassie brilliant blue G at 595 nm. Cryo-milled yeast samples were dissolved in 5% SDS and 50 mM TEAB buffer and digested following the S-Trap mini spin column digestion protocol. Trypsin digested samples were dried in a SpeedVac and resuspended with 20 μL of 0.1% formic acid in water (pH = 3). 2 μL (~ 1 μg digested protein) was injected per run using an Easy-nLC 1200 UPLC system. Samples were loaded directly onto a nano capillary column (45 cm length, 75 μm inner diameter) packed with 1.9 μm C18-AQ resin (Dr. Maisch) mated to metal emitter in-line with an Orbitrap Fusion Lumos (Thermo Fisher Scientific). Column temperature was set at 45°C and two-hour gradient method with 360 nL/min flow was used. The mass spectrometer was operated in data dependent mode with the 120,000 resolution MS1 scan (positive mode, profile data type, AGC gain of 4e5, maximum injection time of 54 sec, and mass range of 375-1500 m/z) in the Orbitrap followed by HCD fragmentation in ion trap with 35% collision energy. Dynamic exclusion list was invoked to exclude previously sequenced peptides for 60 sec and maximum cycle time of 3 sec was used. Peptides were isolated for fragmentation using quadrupole (1.2 m/z isolation window). Ion-trap was operated in Rapid mode. Post-analysis was performed using the Proteome Discoverer v2.5 (Thermo Fisher Scientific). Heterologous protein sequences were added to *S. cerevisiae* CEN.PK113-7D reference database before searching.

### Recombinant Erg20p Expression and Purification

Erg20p wild type and variants were purified using N-terminal 6xHis tag and metal affinity chromatography. Production of the recombinant Erg20p variants was carried out in *E. coli* BL21(DE3) cells harboring the construct plasmids grown overnight at 30°C in 10 mL of Luria-Bertani (LB) broth supplemented with Kanamycin (Kan) at 50 μg/mL. A 1/1000 dilution of the overnight culture was used to inoculate 1 L of LB supplemented with Kan. Protein production was induced by the addition of 0.5 mM IPTG when cell densities were at OD_600_ 0.4~0.8 after incubation at 37°C. After induction, cells were grown at 19 °C for an additional 20 hours, harvested by centrifugation at 7,500 x g for 20 minutes at 4 °C, and resuspended in washing buffer (Tris-buffered saline-TBS, pH 7.6, 10 mL/L culture). To disrupt the cells, the resuspended pellets were then flash-frozen in liquid nitrogen and lysed using the CryoMill system as above. Lysed cells were thawed on ice with the addition of binding buffer (100 mM Tris pH 8.0, 150 mM NaCl, 5 mM imidazole, and 5% v/v glycerol) supplemented with 1 mM PMSF, 2 mg/mL of DNase and 100 µL of commercial protease inhibitor cocktail up to 5% of the initial cell culture volume. Bacterial lysates were clarified by centrifugation at 12,000 x g for 30 minutes at 4°C, then loaded at 1.8 mL/min onto columns of 2.5 mL (50% suspension) of TALON^®^ Superflow™ histidine-tagged protein purification resin previously equilibrated with binding buffer, using a Minipuls^®^ 3 peristaltic pump (Gilson^®^) at 4ºC. Columns were washed with 40–50 column volumes of binding buffer with increasing concentrations of imidazole (5 mM, 10 mM, 20 mM) and proteins eluted with elution buffer (250 mM of imidazole).

At each step of the purification process, 30 µl aliquots were collected for sodium dodecyl sulfate– polyacrylamide gel electrophoresis (SDS–PAGE) analysis. To remove the high concentration of imidazole present in the elution buffer, protein was dialyzed using a Spectra/Por 1 Dialysis Membrane with a molecular weight cutoff of 6-8 kD. The eluted protein was dialyzed in 1 L of dialysis buffer (300 mM NaCl and 20 mM HEPES) for 2 hours and then the buffer was discarded, and 1 L of fresh buffer was added to dialyze the protein for an additional 24hr at 4 °C.

Protein was then concentrated by centrifugation (in cycles of 10 min) at maximum speed in a Sorvall Legend XTR Benchtop centrifuge (Thermo Scientific™) using 15 mL centricons^®^ (Millipore) with a cutoff selected according to the molecular size of each protein, until concentrations between 5 and 10 mg/mL were reached. Finally, the protein was run through Size Exclusion Chromatography (SEC) using a Hiprep™ HR 16/60 Sephacryl™ 200 with Buffer A (50 mM Tris pH 8.0 and 150 mM NaCl) in an FPLC column (ÄKTA pure from GE^®^ Healthcare). The SEC was monitored by UV absorbance at 280 nm. The aliquots enriched with the target proteins were immediately used for the enzymatic assays. The concentrations of the fractions eluted with light were calculated using absorbance at 280 nm with a spectrometer.

### Determination of Kinetic Parameters

Recombinant Erg20p activities were assayed following previous studies with slight modifications^16,73^ by varying the concentrations of DMAPP or GPP added as substrates in a total volume of 0.2 mL reaction containing 10 mM MOPS (pH 7.0), 10 mM MgCl2, 4 mM DTT, 0.1 mg/mL BSA, 0.05 mM IPP, and 50 ng of recombinant Erg20p (wild-type or variants). Reactions were carried out for 30 min at 30°C; and then, they were terminated by the addition of 0.2 mL 2N HCl in 83% ethanol and 0.2 mL ethyl acetate overlay to trap the volatile products. After 10 min at 37°C to hydrolyze the acid-labile diphosphates, reactions were neutralized by adding 0.35 mL 10% NaOH. The ethyl acetate phase (50 μL) was analyzed by GC-FID. The GC method is as follows. Samples were injected and subjected to a split (0.5 μL injection volume; 1:20 split) with a constant flow of 1.5 mL/min. The temperatures of the injector and detector are 230°C and 260°C, respectively. The initial oven temperature was 45°C, increasing at a rate of 1.5°C/min until 150°C, then at 40°C/min until 220°C, being held at this temperature for the last 10 min. For the quantification of products, calibration curves of geraniol, linalool, farnesol, and nerolidol were prepared and diluted in the same sample matrix used for enzyme assays. Enzyme activity was calculated as the total amount of GPP products (geraniol and linalool) or FPP products (farnesol and nerolidol) produced per minute per ng of protein. All the experiments were carried out in triplicate. Graphs and statistical analysis for enzyme kinetics were done using GraphPad PRISM software V10.3.

### Modeling of wild type Erg20p and Erg20p-peptide fusions

ColabFold v1.5.5 with AlphaFold 2.0 parameters was used to produce all structural models^74,75^. The crystal structure of homologous protein isoprenyl diphosphate synthase from *Phaedon cochleariae* (PDB: 8A6V) was provided as a template; the same structure was used to provide an initial pose for IPP in the proximal homoallylic site (HAl) site. DMAPP was positioned in the allylic site (Al) by superimposing the solved DMAPP within the homologous protein structure of isoprenoid synthase A3MSH1 (PDB: 4GP2). GPP was positioned similarly in the Al from the solved pose within homologous protein structure (PDB: 8A70).

The wild-type Erg20p with IPP and GPP or DMAPP posed was subject to energy minimization with the Rosetta *FastRelax* protocol using the “ref2015” score function^76,77^. Optimization of the ligand positioning and pocket sidechains was performed with RosettaLigand using the ROSIE server^78,79^. 200 conformers of each ligand were generated using BCL^80^. Independent trajectories were started from 200 random seeds, and the lowest energy pose for GPP/IPP and DMAPP/IPP was used as the starting pose for generating mutant models.

Secondary structures of a library of C-terminal extensions to Erg20p were determined by RaptorX^81^. Selected mutations and extensions were chosen to build structural models using PyRosetta’s point mutation function ^66^. A constrained *FastRelax* protocol was performed on the additional residues and any residues within a 5Å radius of the extensions. Pocket volume was measured through Fpocket^82,83^. All visualizations were produced with PyMOL (PyMOL Molecular Graphics System, Version 3.0 Schrödinger, LLC).

### Fed-Batch Fermentation in Bioreactor

To test the production of geraniol in large-scale laboratory fermenters, we scaled fermentations up to a 2L bioreactor (Infors HT) using the BioFlo120 system (Eppendorf, B120110001), adding 600 mL SC media supplemented with 2% glucose and 100 μL of Antifoam 204 (Sigma-Aldrich). The reactor was set to maintain a temperature of 30°C with a pH of 5.0, achieved through a 2M ammonium hydroxide supply. The dissolved oxygen levels in the reactor were set to a minimum of 30% through a PID controller that adjusts agitation speed (60-600 rpm) and air flow rate (0.2-2 L/min). The reactor was inoculated with an overnight culture of yKX786 so that the initial OD in the bioreactor was around 0.2. Cell growth was monitored using the BE2100 Biomass Monitor, a noninvasive optical density reader. For in situ extraction of the products, a 10% overlay of isopropyl myristate (IPM) was added. At 24, 48, 72, 96 and 120 hours, the culture was fed with 30 mL of 40% glucose. Timepoints were manually taken, and products (glucose, glycerol, ethanol and geraniol) were quantified through GC-MS and HPLC. Concentrations of glucose, ethanol, and glycerol were measured using HPLC and geraniol was measured using GC-MS. For HPLC measurements, 1mL of aqueous sample was collected and centrifuged in 1.5 mL Eppendorf tubes at 12,500 g for 30 min at 4 °C to remove residual debris. Subsequently, 500 μL of the upper layer was transferred to a 2 mL glass vial for HPLC analysis using an Agilent 6100 system equipped with a BioRad Aminex HPX-87H column (300 × 7.8 mm, 9 μm particle size, 0.25 mm diameter) and a refractive index detector. Both the column and detector temperatures were set to 55°C. Samples were eluted with 5 mM H_2_SO_4_ at a flow rate of 0.6 mL/min for 50 min. To quantify GOH production, 50 μL of the IPM layer was sampled and diluted with 500 μL of ethyl acetate (Sigma Aldrich). The mixture was vortexed for 10 min and then centrifuged at 12,500 g for 20 min at 4°C to remove any residual debris. Then, 450 μL of the supernatant was collected for GC-FID analysis using an Agilent 7890B GC System equipped with a DB-Wax column (30 m length, 0.25 mm diameter, 0.5 μm film). The GC method is described in detail in the Analytical Methods section.

### Analytical Methods

To quantify monoterpenes and farnesene production, 200 μL of the dodecane layer was sampled and diluted with 800 μL of ethyl acetate (Sigma Aldrich). The mixture was vortexed for 5 min and then centrifuged at 12,500 g for 30 min at 4°C to remove any residual debris. Then, 450 μL of the supernatant was collected for GC-FID analysis using an Agilent 7890B GC System equipped with a DB-Wax column (30 m length, 0.25 mm diameter, 0.5 μm film). The GC method is as follows. Samples were injected and subjected to a split injection (0.5 μL injection volume; 1:20 split) with a constant flow of 1.5 mL/min. The oven temperature was initially set to 70°C and held for 5 minutes, followed by a temperature increase at a rate of 30°C/min until reaching 230°C, and then held for 3 minutes. Quantification of the samples was performed using flame-ionization detection (300°C with H_2_ flow at 30 mL/min, air flow at 400 mL/min, and makeup flow at 25 mL/min). Monoterpenes and *β*-farnesene concentrations were calculated using calibration curves generated from standard solutions of geraniol (Sigma-Aldrich), geranyl acetate (Sigma-Aldrich), linalool (Sigma-Aldrich), linalyl Acetate (TCI Chemicals), and *β*-farnesene (a gift from Derek McPhee and Amyris).

Concentrations of glucose, ethanol, and glycerol were measured using HPLC. 1mL of aqueous sample was collected and centrifuged in 1.5 mL Eppendorf tubes at 12,500 g for 30 min at 4 °C to remove residual debris. Subsequently, 500 μL of the upper layer was transferred to a 2 mL glass vial for HPLC analysis using an Agilent 6100 system equipped with a BioRad Aminex HPX-87H column (300 × 7.8 mm, 9 μm particle size, 0.25 mm diameter) and a refractive index detector. Both the column and detector temperatures were set to 55°C. Samples were eluted with 5 mM H_2_SO_4_ at a flow rate of 0.6 mL/min for 50 min.

### Statistical Methods

Statistics for comparing production were calculated using the two-sided t-test. Statistical significance is demoted as: ns (P > 0.05), ^*^ (P < 0.05), ^**^ (P < 0.01), ^***^ (P < 0.001), ^****^ (P < 0.0001).

## References

1. Kamatou, G. P. P. & Viljoen, A. M. Linalool – a Review of a Biologically Active Compound of Commercial Importance. Nat Prod Commun 3, 1934578 0800300 (2008).

2. Wu, T., Li, S., Zhang, B., Bi, C. & Zhang, X. Engineering Saccharomyces cerevisiae for the production of the valuable monoterpene ester geranyl acetate. Microb Cell Fact 17, 1–10 (2018).

3. Chen, L. Separation and purification of plant terpenoids from biotransformation. Eng Life Sci 21, 724–738 (2021).

4. Thomas, A. & Bessiere, Y. The Synthesis of Monoterpenes, 1971—1979 - Thomas - 1981 -. in Total Synthesis of Natural Products (John Wiley & Sons, Ltd).

5. Li, Q. et al. Advances in the sustainable biosynthesis of valuable terpenoid flavor compounds and precursors in micro-organisms. Biotechnology for Biofuels and Bioproducts 18, 99 (2025).

6. Bian, G., Deng, Z. & Liu, T. Strategies for terpenoid overproduction and new terpenoid discovery. Curr Opin Biotechnol 48, 234–241 (2017).

7. Wang, C. Microbial platform for terpenoid production: Escherichia coli and Yeast. Front Microbiol 9, 1–8 (2018).

8. Agranoff, B. W., Eggerer, H., Henning, U. & Lynen, F. Biosynthesis of Terpenes. Journal of Biological Chemistry 235, 326–332 (1960).

9. Ecker, F., Vattekkatte, A., Boland, W. & Groll, M. Metal-dependent enzyme symmetry guides the biosynthetic flux of terpene precursors. Nat. Chem. 15, 1188–1195 (2023).

10. Jiang, L., Huang, L., Cai, J., Xu, Z. & Lian, J. Functional expression of eukaryotic cytochrome P450s in yeast. Biotechnol Bioeng 118, 1050–1065 (2021).

11. Brown, S., Clastre, M., Courdavault, V. & O’Connor, S. E. De novo production of the plant-derived alkaloid strictosidine in yeast. in Proceedings of the National Academy of Sciences vol. 112 3205– 3210 (2015).

12. Oswald, M., Fischer, M., Dirninger, N. & Karst, F. Monoterpenoid biosynthesis in Saccharomyces cerevisiae. FEMS Yeast Res 7, 413–421 (2007).

13. Wollam, J. & Antebi, A. Sterol Regulation of Metabolism, Homeostasis, and Development. Annual Review of Biochemistry 80, 885–916 (2011).

14. Fischer, M. J. C., Meyer, S., Claudel, P., Bergdoll, M. & Karst, F. Metabolic engineering of monoterpene synthesis in yeast. Biotechnol Bioeng 108, 1883–1892 (2011).

15. Zhao, J., Bao, X., Li, C., Shen, Y. & Hou, J. Improving monoterpene geraniol production through geranyl diphosphate synthesis regulation in Saccharomyces cerevisiae. Appl Microbiol Biotechnol 100, 4561–4571 (2016).

16. Ignea, C., Pontini, M., Maffei, M. E., Makris, A. M. & Kampranis, S. C. Engineering monoterpene production in yeast using a synthetic dominant negative geranyl diphosphate synthase. ACS Synth Biol 3, 298–306 (2014).

17. Bernard, A., Cha, S., Shin, H., Lee, D. & Hahn, J.-S. Efficient production of (S)-limonene and geraniol in Saccharomyces cerevisiae through the utilization of an Erg20 mutant with enhanced GPP accumulation capability. Metab Eng 83, 183–192 (2024).

18. Peralta-Yahya, P. P. Identification and microbial production of a terpene-based advanced biofuel. Nat Commun 2, 483–488 (2011).

19. Meadows, A. L. Rewriting yeast central carbon metabolism for industrial isoprenoid production. Nature 537, 694–697 (2016).

20. Paddon, C. J. High-level semi-synthetic production of the potent antimalarial artemisinin. Nature 496, 528–532 (2013).

21. Fischer, M. J. C., Meyer, S., Claudel, P., Bergdoll, M. & Karst, F. Identification of a lysine residue important for the catalytic activity of yeast farnesyl diphosphate synthase. Protein Journal 30, 334– 339 (2011).

22. Lee, H., DeLoache, W. C. & Dueber, J. E. Spatial organization of enzymes for metabolic engineering. Metab Eng 14, 242–251 (2012).

23. Zhao, H., French, J. B., Fang, Y. & Benkovic, S. J. The purinosome, a multi-protein complex involved in the de novo biosynthesis of purines in humans. Chemical Communications 49, 4444 (2013).

24. Jin, M. Glycolytic Enzymes Coalesce in G Bodies under Hypoxic Stress. Cell Rep 20, 895–908 (2017).

25. Kohnhorst, C. L. Identification of a multienzyme complex for glucose metabolism in living cells. Journal of Biological Chemistry 292, 9191–9203 (2017).

26. Walls, M. T., Xu, K., Brangwynne, C. P. & Avalos, J. L. A Modular Design for Synthetic Membraneless Organelles Enables Compositional and Functional Control. 2023.10.03.560789 Preprint at 10.1101/2023.10.03.560789 (2023).

27. Bracha, D. Mapping Local and Global Liquid Phase Behavior in Living Cells Using Photo-Oligomerizable Seeds. Cell 175, 1467–1480 (2018).

28. Rana, U. Asymmetric oligomerization state and sequence patterning can tune multiphase condensate miscibility. Nat Chem https://doi.org/10.1038/s41557-024-01456-6. (2024) doi:10.1038/s41557-024-01456-6.

29. Bellapadrona, G. & Elbaum, M. Supramolecular Protein Assemblies in the Nucleus of Human Cells. Angewandte Chemie International Edition 53, 1534–1537 (2014).

30. Hsia, Y. Design of a hyperstable 60-subunit protein icosahedron. Nature 535, 136–139 (2016).

31. Harris, B. Z., Hillier, B. J. & Lim, W. A. Energetic Determinants of Internal Motif Recognition by PDZ Domains. Biochemistry 40, 5921–5930 (2001).

32. Wegner, S. A. Engineering acetyl-CoA supply and ERG9 repression to enhance mevalonate production in Saccharomyces cerevisiae. J Ind Microbiol Biotechnol 48, (2021).

33. Zhao, L., Xu, L., Westfall, P. & Main, A. Methods of developing terpene synthase variants. (2012).

34. Zhou, P. Combinatorial Modulation of Linalool Synthase and Farnesyl Diphosphate Synthase for Linalool Overproduction in Saccharomyces cerevisiae. J Agric Food Chem 69, 1003–1010 (2021).

35. Dimster-Denk, D. et al. Comprehensive evaluation of isoprenoid biosynthesis regulation in Saccharomyces cerevisiae utilizing the Genome Reporter Matrix™. Journal of Lipid Research 40, 850–860 (1999).

36. Zhou, P. Combining Protein and Organelle Engineering for Linalool Overproduction in Saccharomyces cerevisiae. J Agric Food Chem 71, 10133–10143 (2023).

37. Beekwilder, M. J., Styles, M. Q., Vos, A. M., Hesselink, T. & Jan Bosch, H. Enzymes and methods for fermentative production of monoterpene esters. (2021).

38. Wang, X. Combined bioderivatization and engineering approach to improve the efficiency of geraniol production. Green Chemistry 24, 864–876 (2022).

39. Chacón, M. G., Marriott, A., Kendrick, E. G., Styles, M. Q. & Leak, D. J. Esterification of geraniol as a strategy for increasing product titre and specificity in engineered Escherichia coli. Microb Cell Fact 18, 1–11 (2019).

40. Wang, X. Genetic and Bioprocess Engineering for the Selective and High-Level Production of Geranyl Acetate in Escherichia coli. ACS Sustain Chem Eng 10, 2881–2889 (2022).

41. Castellana, M. Enzyme clustering accelerates processing of intermediates through metabolic channeling. Nat Biotechnol 32, 1011–1018 (2014).

42. Ignea, C. et al. Improving yeast strains using recyclable integration cassettes, for the production of plant terpenoids. Microb Cell Fact 10, 4 (2011).

43. You, S. Utilization of biodiesel by-product as substrate for high-production of B-farnesene via relatively balanced mevalonate pathway in Escherichia coli. Bioresour Technol 243, 228–236 (2017).

44. George, K. W. Integrated analysis of isopentenyl pyrophosphate (IPP) toxicity in isoprenoid-producing Escherichia coli. Metab Eng 47, 60–72 (2018).

45. Primak, Y. A. et al. Characterization of a Feedback-Resistant Mevalonate Kinase from the Archaeon Methanosarcina mazei. Applied and Environmental Microbiology 77, 7772–7778 (2011).

46. Dusséaux, S., Wajn, W. T., Liu, Y., Ignea, C. & Kampranis, S. C. Transforming yeast peroxisomes into microfactories for the efficient production of high-value isoprenoids. Proceedings of the National Academy of Sciences 117, 31789–31799 (2020).

47. Terpe, K. Overview of tag protein fusions: from molecular and biochemical fundamentals to commercial systems. Appl Microbiol Biotechnol 60, 523–533 (2003).

48. Munir, A. Enhanced soluble expression of active recombinant human interleukin-29 using champion pET SUMO system. Biotechnol Lett 45, 1001–1011 (2023).

49. Butt, T. R., Edavettal, S. C., Hall, J. P. & Mattern, M. R. SUMO fusion technology for difficult-to-express proteins. Protein Expr Purif 43, 1–9 (2005).

50. Schoonen, L. et al. Alternative application of an affinity purification tag: hexahistidines in ester hydrolysis. Sci Rep 7, 14772 (2017).

51. Cheah, L. C. Metabolic flux enhancement from the translational fusion of terpene synthases is linked to terpene synthase accumulation. Metab Eng 77, 143–151 (2023).

52. O’Quin, J. B., Mullen, R. T. & Dyer, J. M. Addition of an N-terminal epitope tag significantly increases the activity of plant fatty acid desaturases expressed in yeast cells. Appl Microbiol Biotechnol 83, 117–125 (2009).

53. Zhao, E. M. Light-based control of metabolic flux through assembly of synthetic organelles. Nat Chem Biol 15, 589–597 (2019).

54. Guo, H. Spatial engineering of E. coli with addressable phase-separated RNAs. Cell 185, 3823–3837 23 (2022).

55. Wang, Y. Phase-Separated Multienzyme Compartmentalization for Terpene Biosynthesis in a Prokaryote. Angewandte Chemie International Edition 61, (2022).

56. Zhou, P. Engineered Artificial Membraneless Organelles in Saccharomyces cerevisiae To Enhance Chemical Production. Angewandte Chemie International Edition 62, (2023).

57. Gutensohn, M., Hartzell, E. & Dudareva, N. Another level of complex-ity: The role of metabolic channeling and metabolons in plant terpenoid metabolism. Front. Plant Sci. 13, (2022).

58. Camagna, M. et al. Enzyme Fusion Removes Competition for Geranylgeranyl Diphosphate in Carotenogenesis. Plant Physiol 179, 1013–1027 (2019).

59. Zhang, Y. & Fernie, A. R. Metabolons, enzyme–enzyme assemblies that mediate substrate channeling, and their roles in plant metabolism. Plant Communications 2, 100081 (2021).

60. Wei, S.-P. et al. Formation and functionalization of membraneless compartments in Escherichia coli. Nat Chem Biol 16, 1143–1148 (2020).

61. Hilditch, A. T. et al. Assembling membraneless organelles from de novo designed proteins. Nat. Chem. 16, 89–97 (2024).

62. Yu, W. et al. De novo engineering of programmable and multi-functional biomolecular condensates for controlled biosynthesis. Nat Commun 15, 7989 (2024).

63. Gil-Garcia, M. et al. Local environment in biomolecular condensates modulates enzymatic activity across length scales. Nat Commun 15, 3322 (2024).

64. Stoffel, F. et al. Enhancement of enzymatic activity by biomolecular condensates through pH buffering. Nat Commun 16, 6368 (2025).

65. Peeples, W. & Rosen, M. K. Mechanistic dissection of increased enzymatic rate in a phase-separated compartment. Nat Chem Biol 17, 693–702 (2021).

66. Giannakopoulou, A. et al. Design of biomolecular condensates for sustainable bioproduction of acetoin. In preparation.

67. Matsuzawa, T. et al. Metabolites Shift Equilibria of Biomolecular Condensates. 2026.01.14.699531 Preprint at 10.64898/2026.01.14.699531 (2026).

68. Siu, K.-H. et al. Spatial organization of enzymes and cofactors in synthetic membrane-associated condensates (sMACs). Submitted.

69. Stovicek, V., Borja, G. M., Forster, J. & Borodina, I. EasyClone 2.0: expanded toolkit of integrative vectors for stable gene expression in industrial Saccharomyces cerevisiae strains. J Ind Microbiol Biotechnol 42, 1519–1531 (2015).

70. Lee, M. E., DeLoache, W. C., Cervantes, B. & Dueber, J. E. A Highly Characterized Yeast Toolkit for Modular, Multipart Assembly. ACS Synth Biol 4, 975–986 (2015).

71. Gietz, R. D. & Woods, R. A. Transformation of yeast by lithium acetate/single-stranded carrier DNA/polyethylene glycol method. Methods Enzymol 350, 87–96 (2002).

72. Shaw, W. M. Engineering a Model Cell for Rational Tuning of GPCR Signaling. Cell 177, 782–796 27 (2019).

73. Ferriols, V. M. E. N. Cloning and characterization of farnesyl pyrophosphate synthase from the highly branched isoprenoid producing diatom Rhizosolenia setigera. Sci Rep 5, 10246 (2015).

74. Jumper, J. Highly accurate protein structure prediction with AlphaFold. Nature 596, 583–589 (2021).

75. Mirdita, M. ColabFold: making protein folding accessible to all. Nat Methods 19, 679–682 (2022).

76. Chaudhury, S., Lyskov, S. & Gray, J. J. PyRosetta: a script-based interface for implementing molecular modeling algorithms using Rosetta. Bioinformatics 26, 689–691 (2010).

77. Conway, P., Tyka, M. D., DiMaio, F., Konerding, D. E. & Baker, D. Relaxation of backbone bond geometry improves protein energy landscape modeling. Protein Science 23, 47–55 (2014).

78. DeLuca, S., Khar, K. & Meiler, J. Fully Flexible Docking of Medium Sized Ligand Libraries with RosettaLigand. PLOS ONE 10, e0132508 (2015).

79. Lyskov, S. Serverification of Molecular Modeling Applications: The Rosetta Online Server That Includes Everyone (ROSIE. PLoS One 8, e63906, (2013).

80. Kothiwale, S., Mendenhall, J. L. & Meiler, J. BCL::Conf: small molecule conformational sampling using a knowledge based rotamer library. Journal of Cheminformatics 7, 47 (2015).

81. Wang, S., Li, W., Liu, S. & Xu, J. RaptorX-Property: a web server for protein structure property prediction. Nucleic Acids Research 44, W430–W435 (2016).

82. Le Guilloux, V., Schmidtke, P. & Tuffery, P. Fpocket: An open source platform for ligand pocket detection. BMC Bioinformatics 10, 168 (2009).

83. Kochnev, Y. & Durrant, J. D. FPocketWeb: protein pocket hunting in a web browser. Journal of Cheminformatics 14, 58 (2022).

