## Supplemental Information for "Enhanced GPP synthesis by Erg20p-peptide fusions and biomolecular condensates boosts monoterpene production in yeast"

### C-terminus peptide fusion to Erg20p enhances its GPP synthase activity and enables spatial organization of enzymes to improve monoterpene production in *Saccharomyces cerevisiae*

Ke Xu[1*], Archontoula Giannakopoulou[1*], Virginia Jiang[1*], Saurabh Malani[1], Mackenzie T. Walls[1], Clifford P. Brangwynne[1,2,3], José L. Avalos[1,3,4]

1. Department of Chemical and Biological Engineering, Princeton University, Princeton NJ USA
2. Howard Hughes Medical Institute, Chevy Chase, MD USA
3. Omenn-Darling Bioengineering Institute, Princeton University, Princeton, NJ, USA
4. Andlinger Center for Energy and the Environment, Princeton University, Princeton NJ USA
* Co-First Authors

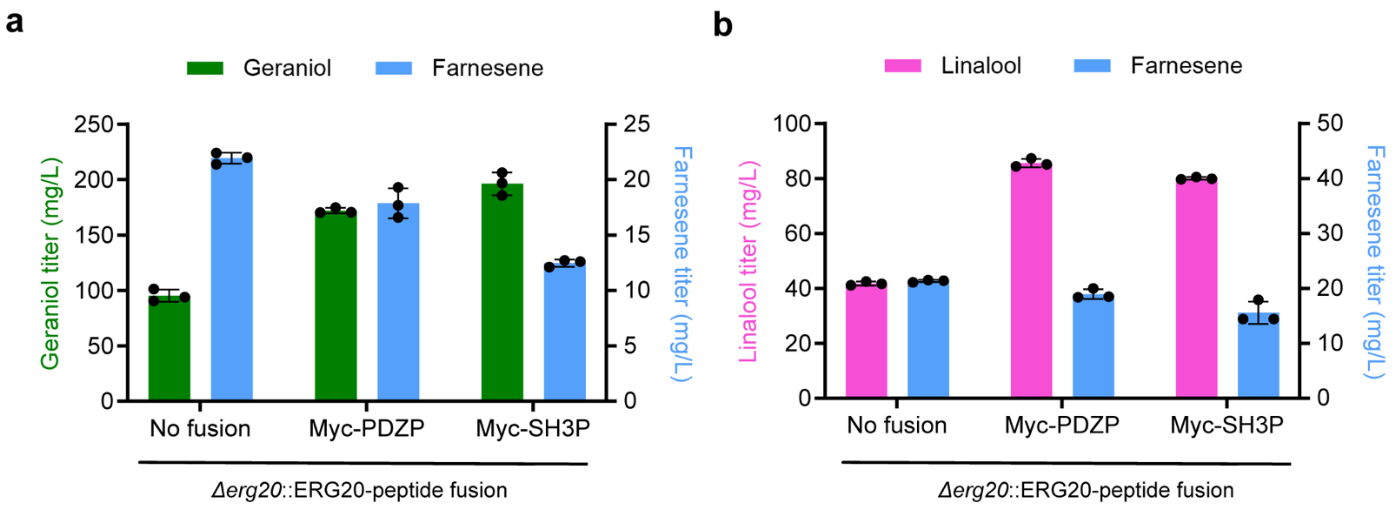


**Figure S1: Monoterpenes and farnesene production in yeast strains expressing Erg20p fused with C-terminus Myc–PDZ$_P$ and Myc–SH3$_P$ peptides**. **(a)** Geraniol production and **(b)** Linalool production. Error bars represent one standard deviation from three biologically replicates.

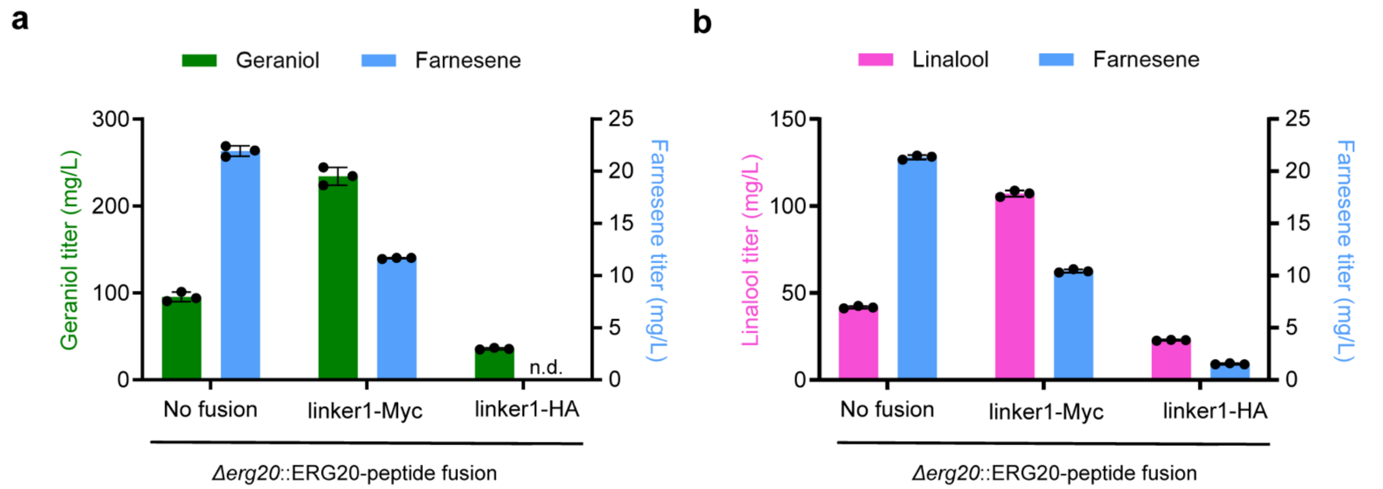


**Figure S2: Monoterpenes and farnesene production in yeast strains expressing Erg20p fused with C-terminus linker1–Myc and linker1–HA peptides**. **(a)** Geraniol production and **(b)** Linalool production. Error bars represent one standard deviation from three biologically replicates.

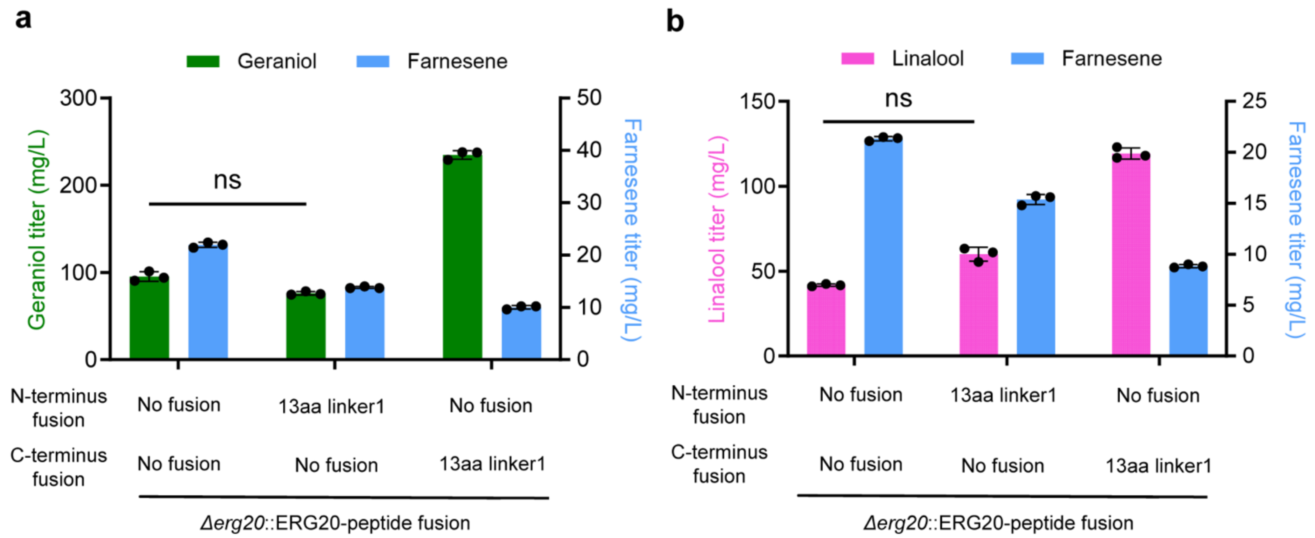


**Figure S3: Monoterpenes and farnesene production in yeast strains expressing Erg20p fused with either N-terminus or C-terminus 13aa linker1 peptides**. **(a)** Geraniol production and **(b)** Linalool production. Error bars represent one standard deviation from three biologically replicates.

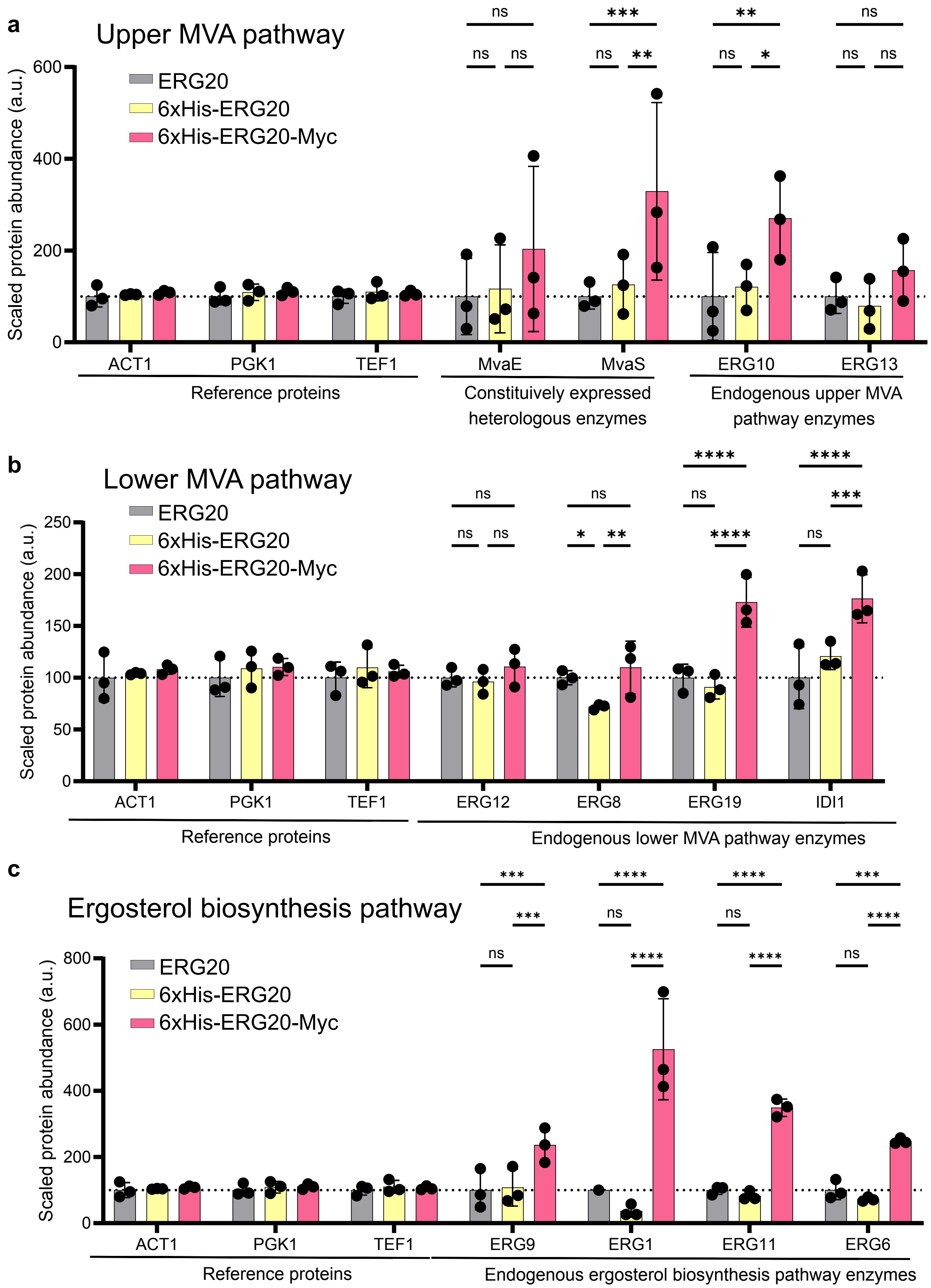


**Figure S4: The relative abundances of native protein involved in the biosynthesis of ergosterol. (a)** The upper mevalonate pathway. **(b)** The lower mevalonate pathway. **(c)** Ergosterol biosynthesis pathway. Error bars represent one standard deviation from three biologically replicates. Values are normalized so that the abundance of the given protein in the wild-type *ERG20* strain is equal to 100. Statistics are calculated using a two-sided t-test. ns (non-significant); *P < 0.05; ***P < 0.01; ***P < 0.001; ***P < 0.0001.

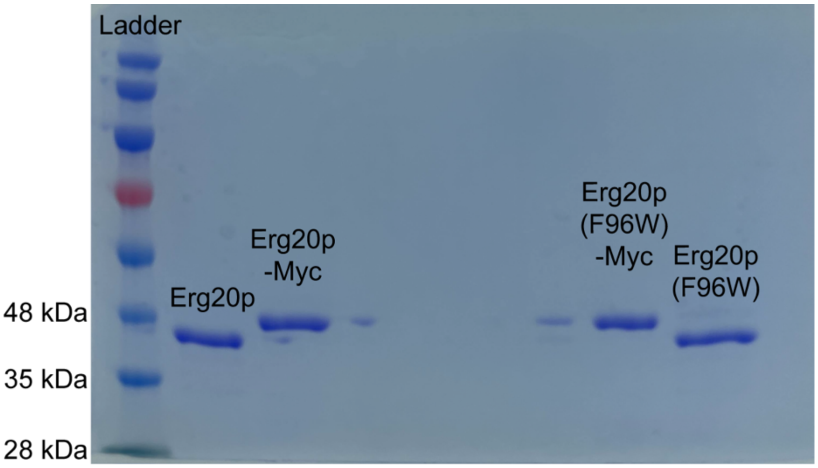


**Figure S5: SDS-PAGE gel of the recombinant proteins purified from *E. coli*.** All proteins are fused to a N-terminal 6xHis tag and purified using immobilized metal affinity chromatography.

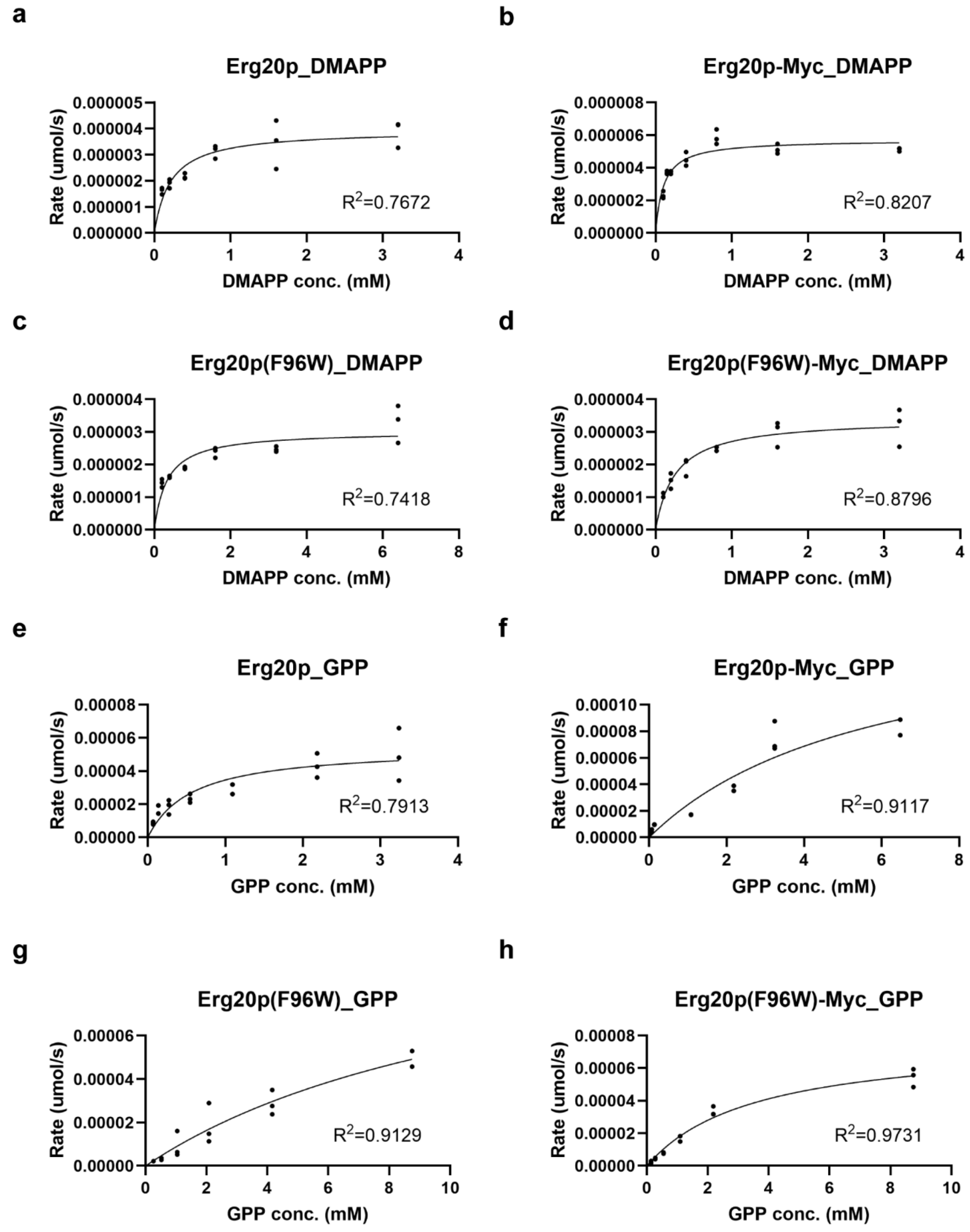


**Figure S6: Steady-state kinetics of Erg20p wild-type and variants. (a-d)** DMAPP as substrate. **(e-f)** GPP as substrate. Data fitting to Michaelis–Menten kinetics is performed in GraphPad PRISM software.

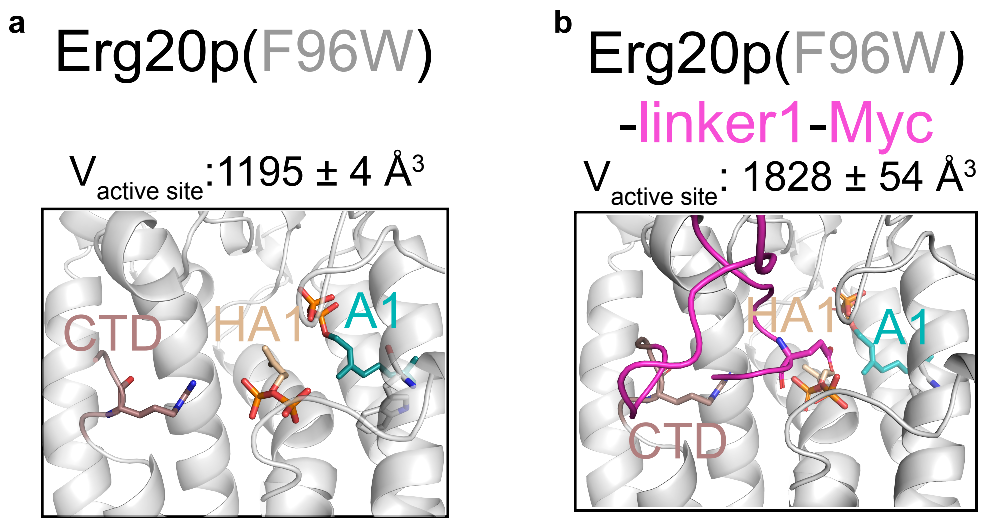


**Figure S7: Impact of C-terminal fusion and codon 96 mutation on relative catalytic efficiency for GPP synthase activity or FPP synthase activity. (a)** Structural model of Erg20p(F96W) and **(b)** Erg20p(F96W)-linker1-Myc fusion, including active-site pocket volume calculations. The side chain of residue 96 is shown as gray sticks.

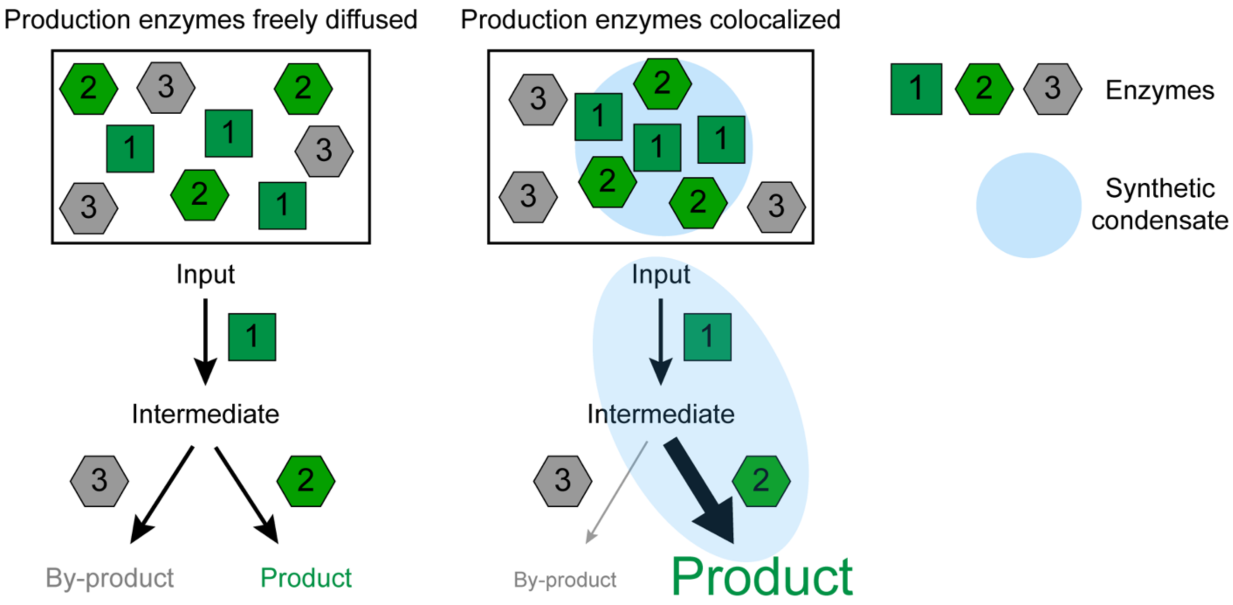


**Figure S8: Synthetic condensates can act as a scaffold to recruit metabolic pathway enzymes for regulating metabolic flux.** When pathway enzymes are colocalized, the flux of an intermediate towards the desired product is enhanced over the formation of an unwanted by-product.

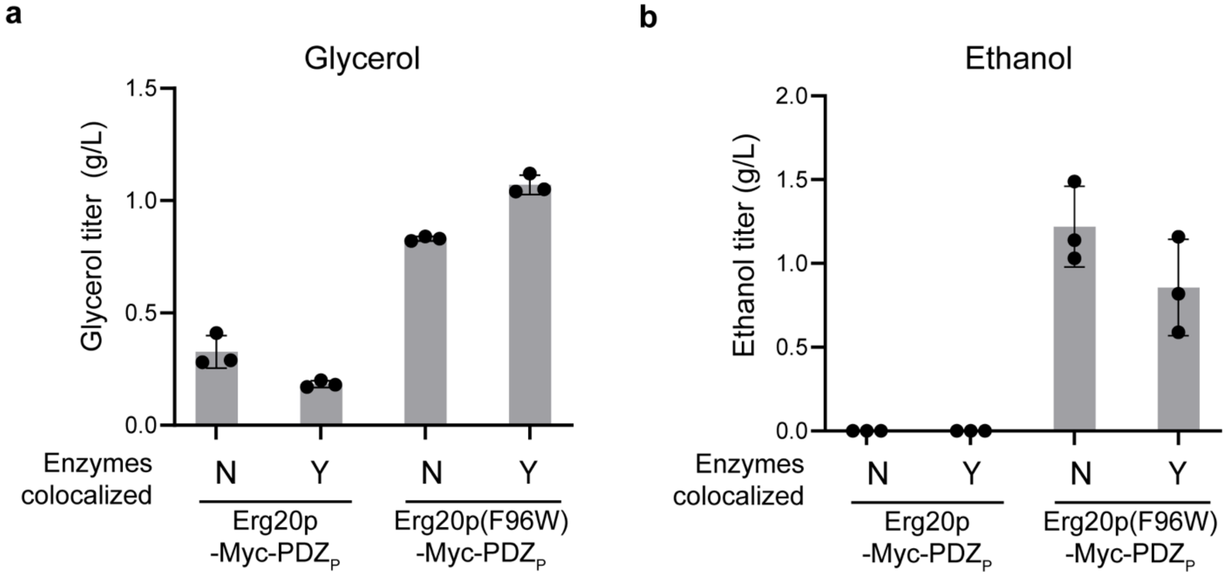


**Figure S9: Accumulation of overflow metabolites in yeast strains producing geraniol with and without Erg20p–Myc–PDZ$_P$ and ObGES–PDZ$_P$ colocalized, and with or without the F96W mutation in Erg20p.** **(a)** Glycerol. **(b)** Ethanol. Error bars represent one standard deviation from three biologically replicates.

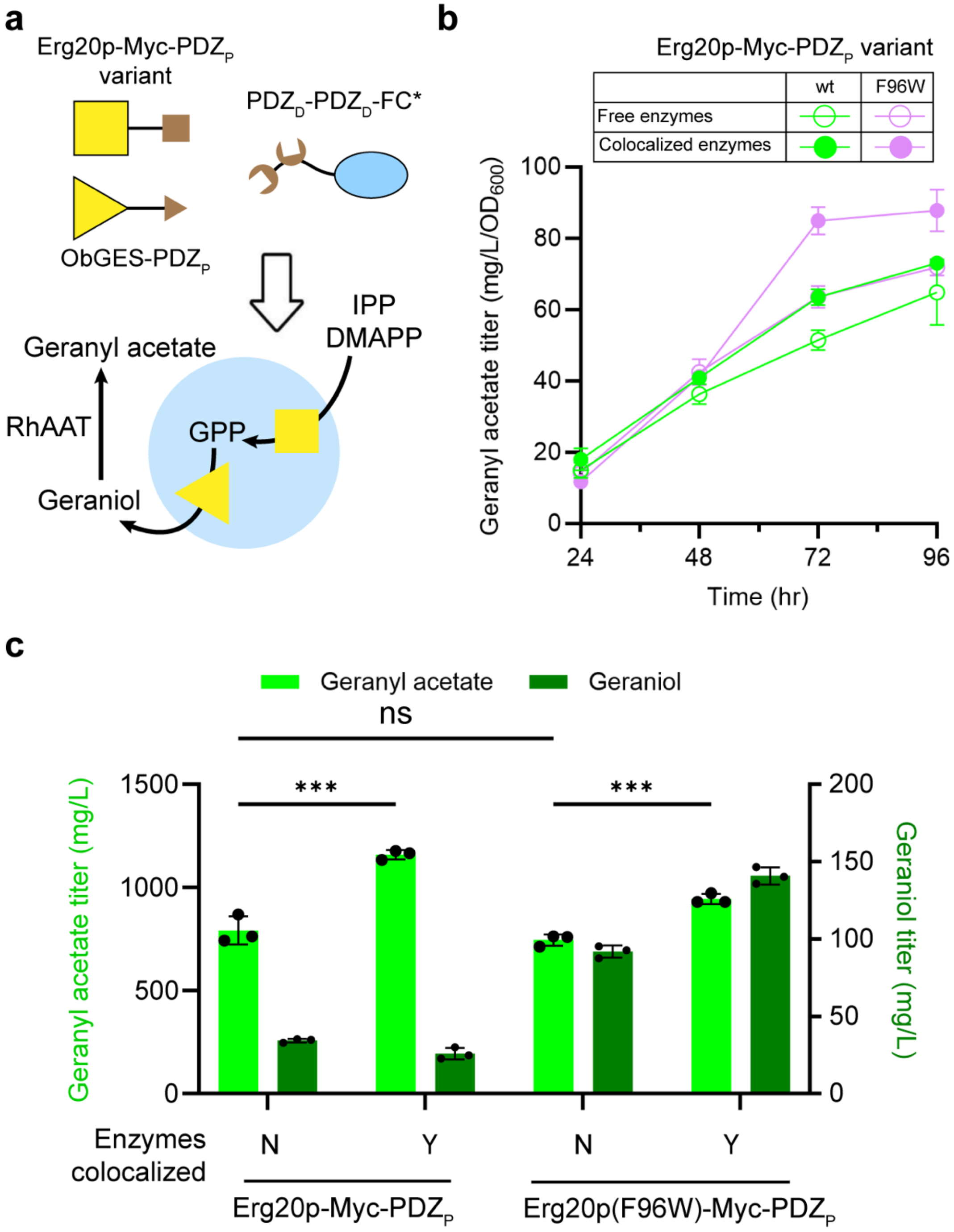


**Figure S10: Improve geranyl acetate production via synthetic condensate-mediated colocalization of Erg20p and geraniol synthase. (a)** Schematics of colocalizing Erg20p and ObGES into Forever Corelet (FC*) synthetic condensates using protein domain-peptide interactive pairs to streamline geraniol production. RhAAT is overexpressed to convert geraniol to geranyl acetate. **(b)** Time-course of $OD_{600}$-normalized geranyl acetate production with and without Erg20p–Myc–$PDZ_P$ and ObGES–$PDZ_P$ colocalized, and with or without the F96W mutation in Erg20p. **(c)** The final geranyl acetate titers of yeast strains in (b) after 96hr fermentation. Error bars represent one standard deviation from three biologically replicates. Statistics are calculated using a two-sided t-test. ns (non-significant); ***$P < 0.001$.

**Supplementary Table 1. Peptide primary sequences**

Where present, epitope tags are indicated by underlining

| Sequence | Description |
|---|---|
| linker1–Myc | LEGGSAAGTGSGGEQKLISEEDL |
| linker1–HA | LEGGSAAGTGSGGYPYDVPDYA |
| 13aa linker1 | LEGGSAAGTGSGG |
| linker2 | VEGGGGSGGGGS |
| 7aa linker1 | LEGGSAA |
| 2aa linker1 | LE |

**Supplementary Table 2. Plasmids used in this study.**

| Plasmid | Description | Source |
|---|---|---|
| pKX683 | Amp$^R$, HIS3 integration, TEF1p_AaFSmut-Myc_SSA1t_<br>Rev(TDH3p_AaFSmut-Myc_ACT1t) | This study |
| pKX628 | Amp$^R$, LEU2 integration, CCW12p_ObGES-Myc-PDZ$_P$_ENO1t_<br>Rev(HHF2p_ObGES-HA-PDZ$_P$_ADH1t) | This study |
| pKX577 | Amp$^R$, LEU2 integration, CCW12p_SKIK-McLIS(E343D-E352H)-Myc-PDZ$_P$_ENO1t_<br>Rev(HHF2p_SKIK-McLIS(E343D-E352H)-HA-PDZ$_P$_ADH1t) | This study |
| pKX624 | Amp$^R$, LEU2 integration, PGK1p_coERG20(S288C)-Myc_ACT1t_<br>CCW12p_ObGES-Myc-PDZ$_P$_ENO1t_<br>Rev(HHF2p_ObGES-HA-PDZ$_P$_ADH1t) | This study |
| pKX761 | Amp$^R$, LEU2 integration, PGK1p_coERG20(S288C)-Myc_ACT1t_<br>CCW12p_SKIK-McLIS(E343D-E352H)-Myc-PDZ$_P$_ENO1t_<br>Rev(HHF2p_SKIK-McLIS(E343D-E352H)-HA-PDZ$_P$_ADH1t) | This study |
| pKX554 | Amp$^R$, HIS3 integration, CCW12p_ObGES-Myc-PDZ$_P$_ENO1t_<br>Rev(PGK1p_ObGES-Myc-PDZ$_P$_ACT1t) | This study |
| pKX678 | Amp$^R$, XII-2 Integration (*KanMX*), TEF1p_ObGES-Myc-PDZ$_P$_PGK1t_<br>Rev(PGK1p_ObGES-Myc-PDZ$_P$_ACT1t) | This study |
| pKX629 | Amp$^R$, LEU2 integration, CCW12p_ObGES-Myc-PDZ$_P$_ENO1t_<br>Rev(TDH3p_RhAAT-Myc_PGK1t)_<br>Rev(HHF2p_ObGES-HA-PDZ$_P$_ADH1t) | This study |
| pKX583 | Amp$^R$, HIS3 integration, CCW12p_SKIK-McLIS(E343D-E352H)-Myc-PDZ$_P$_ENO1t_<br>Rev(TDH3p_SKIK-McLIS(E343D-E352H)-HA-PDZ$_P$_PGK1t) | This study |
| pKX704 | Amp$^R$, XII-2 Integration (*KanMX*)<br>CCW12p_SKIK-McLIS(E343D-E352H)-Myc-PDZ$_P$_ENO1t_<br>Rev(PGK1p_SKIK-McLIS(E343D-E352H)-HA-PDZ$_P$_ACT1t) | This study |
| pKX710 | Amp$^R$, LEU2 integration, TEF1p_CbAAT9-1-c-Myc_SSA1t_<br>CCW12p_SKIK-McLIS(E343D-E352H)-Myc-PDZ$_P$_ENO1t_<br>Rev(TDH3p_CbAAT9-1-c-Myc_PGK1t)_<br>Rev(HHF2p_SKIK-McLIS(E343D-E352H)-HA-PDZ$_P$_ADH1t) | This study |

| | | |
|---|---|---|
| pKX712 | Amp$^R$, HIS3 integration, TDH3p_CbAAT9-1-c-Myc_PGK1t_<br>CCW12p_SKIK-McLIS(E343D-E352H)-Myc-PDZ$_P$_ENO1t_<br>Rev(TEF1p_CbAAT9-1-c-Myc_SSA1t)<br>Rev(TDH3p_SKIK-McLIS(E343D-E352H)-HA-PDZ$_P$_PGK1t) | This study |
| pKX714 | Amp$^R$, XII-2 Integration (*KanMX*), TDH3p_CbAAT9-1-c-Myc_PGK1t_<br>CCW12p_SKIK-McLIS(E343D-E352H)-Myc-PDZ$_P$_ENO1t_<br>Rev(TEF1p_CbAAT9-1-c-Myc_SSA1t)<br>Rev(PGK1p_SKIK-McLIS(E343D-E352H)-HA-PDZ$_P$_ACT1t) | This study |
| pKX654 | Amp$^R$, TRP1 integration, TDH3p_ERG19_CYC1t_TEF1p_ERG8_ACT1t_<br>PGK1p_ERG12_ADH1t_CCW12p_IDI1_ENO1t | This study |
| pKX646 | Amp$^R$, LEU2 integration, CCW12p_ObGES-Myc-PDZ$_P$_ENO1t_<br>TDH3p_PDZ-PDZ-FUSn-mCherry-I301(K129A)_ADH1t_<br>Rev(TEF1p_PDZ-PDZ-FUSn-mCherry-I301(K129A)_TDH1t)_<br>Rev(HHF2p_ObGES-HA-PDZ$_P$_ADH1t) | This study |
| pKX555 | Amp$^R$, HIS3 integration,<br>Rev(TDH3p_PDZ-PDZ-FUSn-mCherry-I301(K129A)_ADH1t)_<br>CCW12p_ObGES-Myc-PDZ$_P$_ENO1t_<br>TEF1p_PDZ-PDZ-FUSn-mCherry-I301(K129A)_TDH1t_<br>Rev(PGK1p_ObGES-Myc-PDZ$_P$_ACT1t) | This study |
| pKX674 | Amp$^R$, XII-2 Integration (*KanMX*), TEF1p_ObGES-Myc-PDZP_PGK1t_<br>CCW12p_PDZ-PDZ-FUSn-mCherry-I301(K129A)_ENO1t_<br>Rev(PGK1p_ObGES-Myc-PDZP_ACT1t)_<br>Rev(TDH3p_PDZ-PDZ-FUSn-mCherry-I301(K129A)_TDH1t) | This study |
| pKX648 | Amp$^R$, LEU2 integration, CCW12p_ObGES-Myc-PDZ$_P$_ENO1t_<br>TDH3p_PDZ-PDZ-FUSn-mCherry-I301(K129A)_ADH1t_<br>Rev(TEF1p_PDZ-PDZ-FUSn-mCherry-I301(K129A)_TDH1t)_<br>Rev(TDH3p_RhAAT-Myc_PGK1t)_<br>Rev(HHF2p_ObGES-HA-PDZ$_P$_ADH1t) | This study |
| pKX647 | Amp$^R$, LEU2 integration, CCW12p_ObGES-Myc-PDZ$_P$_ENO1t_<br>TDH3p_FUSn-mCherry-I301(K129A)_ADH1t_<br>Rev(TEF1p_FUSn-mCherry-I301(K129A)_TDH1t)_<br>Rev(HHF2p_ObGES-HA-PDZ$_P$_ADH1t) | This study |
| pKX556 | Amp$^R$, HIS3 integration, Rev(TDH3p_FUSn-mCherry-I301(K129A)_ADH1t)_<br>CCW12p_ObGES-Myc-PDZ$_P$_ENO1t_<br>TEF1p_FUSn-mCherry-I301(K129A)_TDH1t_<br>Rev(PGK1p_ObGES-Myc-PDZ$_P$_ACT1t) | This study |
| pKX676 | Amp$^R$, XII-2 Integration (*KanMX*), TEF1p_ObGES-Myc-PDZ$_P$_PGK1t_<br>CCW12p_FUSn-mCherry-I301(K129A)_ENO1t_<br>Rev(PGK1p_ObGES-Myc-PDZ$_P$_ACT1t)_<br>Rev(TDH3p_FUSn-mCherry-I301(K129A)_TDH1t) | This study |
| pKX649 | Amp$^R$, LEU2 integration, CCW12p_ObGES-Myc-PDZ$_P$_ENO1t_<br>TDH3p_FUSn-mCherry-I301(K129A)_ADH1t_<br>Rev(TEF1p_FUSn-mCherry-I301(K129A)_TDH1t)_<br>Rev(TDH3p_RhAAT-Myc_PGK1t)_<br>Rev(HHF2p_ObGES-HA-PDZ$_P$_ADH1t) | This study |
| pKX173 | Amp$^R$, LEU2 integration, empty plasmid | This study |

| | | |
|---|---|---|
| pYZ163 | $Amp^R$, HIS3 integration, empty plasmid | This study |
| pYZ161 | $Amp^R$, TRP1 integration, empty plasmid | This study |
| pKX758 | $Amp^R$, URA3 integration, empty plasmid | This study |
| pKX557 | $Amp^R$, CRISPR knock-in, ERG20 sgRNA | This study |
| pKX532 | $Amp^R$, CRISPR knock-in, coERG20(S288C)-Myc-$PDZ_P$ | This study |
| pKX300 | $Amp^R$, CRISPR knock-in, coERG20(S288C)-Myc-$SH3_P$ | This study |
| pKX533 | $Amp^R$, CRISPR knock-in, coERG20(F96W)(S288C)-Myc-$PDZ_P$ | This study |
| pKX660 | $Amp^R$, CRISPR knock-in, coERG20(S288C)-Myc | This study |
| pKX718 | $Amp^R$, CRISPR knock-in, coERG20(S288C)-HA | This study |
| pKX717 | $Amp^R$, CRISPR knock-in, coERG20(S288C)-13aa linker1 | This study |
| pKX730 | $Amp^R$, CRISPR knock-in, coERG20(S288C)-7aa linker1 | This study |
| pKX731 | $Amp^R$, CRISPR knock-in, coERG20(S288C)-2aa linker1 | This study |
| pKX757 | $Amp^R$, CRISPR knock-in, 13aa linker1-coERG20(S288C) | This study |
| pKX715 | $Amp^R$, CRISPR knock-in, 6xHis-coERG20(S288C) | This study |
| pKX716 | $Amp^R$, CRISPR knock-in, 6xHis-coERG20(S288C)-Myc | This study |
| pWS082 | $Amp^R$, CRISPR, sgRNA entry vector | [1] |
| pWS172 | $Kan^R$, CRISPR, Cas9 gap repair vector - HIS3 | [1] |
| pWS173 | $Kan^R$, CRISPR, Cas9 gap repair vector - KanR | [1] |
| pAG4 | $Kan^R$, purifying proteins from *E.coli*, $P_{T7}$_6xHis-coERG20(S288C) | This study |
| pAG5 | $Kan^R$, purifying proteins from *E.coli*, $P_{T7}$_6xHis-coERG20(S288C)-Myc | This study |
| pAG6 | $Kan^R$, purifying proteins from *E.coli*, $P_{T7}$_6xHis-coERG20(F96W)(S288C)-Myc | This study |
| pKS922 | $Kan^R$, purifying proteins from *E.coli*, $P_{T7}$_6xHis-coERG20(F96W)(S288C) | This study |

**Supplementary Table 3. Yeast strains used in this study.**

| Strain | Description | Figure | Genotype | Source |
|---|---|---|---|---|
| CEN.PK2-1C | Wild-type *Saccharomyces cerevisiae* | | *MATa ura3-52 trp1-289 leu2-3,112 his3-1 MAL2-8c SUC2* | [2] |
| JCY51 | Mevalonate overproducing strain | | CEN.PK2-1C δ::TDH3p_EfmvaE_ADH1t_ TEF1p_EfmvaS_ACT1t_ PGK1p_SeACS(L641P)_CYC1t | [3] |
| yKX607 | FPP vs. GPP (geraniol producing, no peptide fusion) | Fig. 2b, S1a, S2a, Fig 3a, Fig 3b, S3a, Fig 4d, | JCY51, ΔERG20::coERG20(S288C) *his3::HIS3*-TEF1p_AaFSmut-Myc_SSA1t_ Rev(TDH3p_AaFSmut-Myc_ACT1t) *leu2::LEU2*-CCW12p_ObGES-Myc-$PDZ_P$_ENO1t_ Rev(HHF2p_ObGES-HA-$PDZ_P$_ADH1t) | This study |

| | | | | |
|---|---|---|---|---|
| | | S6a, S6b, S6c | | |
| yKX690 | FPP vs. GPP (geraniol producing, Erg20p-Myc-PDZ$_P$ fusion) | Fig. 2b, S1a, S2b, Fig 6e | JCY51, ΔERG20::coERG20(S288C)-Myc-PDZ$_P$<br>*his3::HIS3*-TEF1p_AaFSmut-Myc_SSA1t_ Rev(TDH3p_AaFSmut-Myc_ACT1t)<br>*leu2::LEU2*-CCW12p_ObGES-Myc-PDZ$_P$_ENO1t_ Rev(HHF2p_ObGES-HA-PDZ$_P$_ADH1t) | This study |
| yKX692 | FPP vs. GPP (geraniol producing, Erg20p-Myc-SH3$_P$ fusion) | S1a | JCY51, ΔERG20::coERG20(S288C)-Myc-SH3$_P$<br>*his3::HIS3*-TEF1p_AaFSmut-Myc_SSA1t_ Rev(TDH3p_AaFSmut-Myc_ACT1t)<br>*leu2::LEU2*-CCW12p_ObGES-Myc-PDZ$_P$_ENO1t_ Rev(HHF2p_ObGES-HA-PDZ$_P$_ADH1t) | This study |
| yKX608 | FPP vs. GPP (geraniol producing, Erg20p-Myc fusion) | Fig 2b | JCY51, ΔERG20::coERG20(S288C)-Myc<br>*his3::HIS3*-TEF1p_AaFSmut-Myc_SSA1t_ Rev(TDH3p_AaFSmut-Myc_ACT1t)<br>*leu2::LEU2*-CCW12p_ObGES-Myc-PDZ$_P$_ENO1t_ Rev(HHF2p_ObGES-HA-PDZ$_P$_ADH1t) | This study |
| yKX721 | FPP vs. GPP (geraniol producing, Erg20p-Myc-HA fusion) | S2a | JCY51, ΔERG20::coERG20(S288C)-HA<br>*his3::HIS3*-TEF1p_AaFSmut-Myc_SSA1t_ Rev(TDH3p_AaFSmut-Myc_ACT1t)<br>*leu2::LEU2*-CCW12p_ObGES-Myc-PDZ$_P$_ENO1t_ Rev(HHF2p_ObGES-HA-PDZ$_P$_ADH1t) | This study |
| yKX720 | FPP vs. GPP (geraniol producing, Erg20p-13 aa linker fusion) | Fig 2b, S3a | JCY51, ΔERG20::coERG20(S288C)-linker1<br>*his3::HIS3*-TEF1p_AaFSmut-Myc_SSA1t_ Rev(TDH3p_AaFSmut-Myc_ACT1t)<br>*leu2::LEU2*-CCW12p_ObGES-Myc-PDZ$_P$_ENO1t_ Rev(HHF2p_ObGES-HA-PDZ$_P$_ADH1t) | This study |
| yKX797 | FPP vs. GPP (geraniol producing, 13 aa linker -Erg20p fusion) | S3a | JCY51, ΔERG20::linker1-coERG20(S288C)<br>*his3::HIS3*-TEF1p_AaFSmut-Myc_SSA1t_ Rev(TDH3p_AaFSmut-Myc_ACT1t)<br>*leu2::LEU2*-CCW12p_ObGES-Myc-PDZ$_P$_ENO1t_ Rev(HHF2p_ObGES-HA-PDZ$_P$_ADH1t) | This study |
| yKX793 | FPP vs. GPP geraniol producing, Erg20p-7 aa linker fusion) | Fig 2b | JCY51, ΔERG20::coERG20(S288C)-7aa linker1<br>*his3::HIS3*- TEF1p_AaFSmut-Myc_SSA1t_ Rev(TDH3p_AaFSmut-Myc_ACT1t)<br>*leu2::LEU2*-CCW12p_ObGES-Myc-PDZ$_P$_ENO1t_ Rev(HHF2p_ObGES-HA-PDZ$_P$_ADH1t) | This study |
| yKX794 | FPP vs. GPP (geraniol producing, Erg20p-2 aa linker fusion LE) | Fig 2b, 4d | JCY51, ΔERG20::coERG20(S288C)-2aa linker1<br>*his3::HIS3*-TEF1p_AaFSmut-Myc_SSA1t_ | This study |

| | | | | |
|---|---|---|---|---|
| | | | Rev(TDH3p_AaFSmut-Myc_ACT1t)<br>*leu2::LEU2*-CCW12p_ObGES-Myc-PDZ$_P$_ENO1t_<br>Rev(HHF2p_ObGES-HA-PDZ$_P$_ADH1t) | |
| yKX718 | FPP vs. GPP (geraniol producing, 6xHis-Erg20p) | Fig 3a, 3b, S6a, S6b, S6c | JCY51, ΔERG20::6xhis-coERG20(S288C)<br>*his3::HIS3*-TEF1p_AaFSmut-Myc_SSA1t_<br>Rev(TDH3p_AaFSmut-Myc_ACT1t)<br>*leu2::LEU2*-CCW12p_ObGES-Myc-PDZ$_P$_ENO1t_<br>Rev(HHF2p_ObGES-HA-PDZ$_P$_ADH1t) | This study |
| yKX719 | FPP vs. GPP (geraniol producing, 6xHis-Erg20p-Myc) | Fig 3a, 3b, S6a, S6b, S6c | JCY51, ΔERG20::6xhis-coERG20(S288C)-Myc<br>*his3::HIS3*-TEF1p_AaFSmut-Myc_SSA1t_<br>Rev(TDH3p_AaFSmut-Myc_ACT1t)<br>*leu2::LEU2*-CCW12p_ObGES-Myc-PDZ$_P$_ENO1t_<br>Rev(HHF2p_ObGES-HA-PDZ$_P$_ADH1t) | This study |
| yKX660 | FPP vs. GPP (linalool producing, no peptide fusion) | Fig 2c, S1b, S3b | JCY51, ΔERG20::coERG20(S288C)<br>*his3::HIS3*-TEF1p_AaFSmut-Myc_SSA1t_<br>Rev(TDH3p_AaFSmut-Myc_ACT1t)<br>*leu2::LEU2*-CCW12p_SKIK-McLIS(E343D-E352H)-Myc-PDZ$_P$_ENO1t_<br>Rev(HHF2p_SKIK-McLIS(E343D-E352H)-HA-PDZ$_P$_ADH1t) | This study |
| yKX730 | FPP vs. GPP (linalool producing, Erg20p-Myc-PDZ$_P$ fusion) | Fig 2c, S1b | JCY51, ΔERG20::coERG20(S288C)-Myc-PDZ$_P$<br>*his3::HIS3*-TEF1p_AaFSmut-Myc_SSA1t_<br>Rev(TDH3p_AaFSmut-Myc_ACT1t)<br>*leu2::LEU2*-CCW12p_SKIK-McLIS(E343D-E352H)-Myc-PDZ$_P$_ENO1t_<br>Rev(HHF2p_SKIK-McLIS(E343D-E352H)-HA-PDZ$_P$_ADH1t) | This study |
| yKX731 | FPP vs. GPP (linalool producing, Erg20p-Myc-SH3$_P$ fusion) | S1b | JCY51, ΔERG20::coERG20(S288C)-Myc-SH3$_P$<br>*his3::HIS3*-TEF1p_AaFSmut-Myc_SSA1t_<br>Rev(TDH3p_AaFSmut-Myc_ACT1t)<br>*leu2::LEU2*-CCW12p_SKIK-McLIS(E343D-E352H)-Myc-PDZ$_P$_ENO1t_<br>Rev(HHF2p_SKIK-McLIS(E343D-E352H)-HA-PDZ$_P$_ADH1t) | This study |
| yKX661 | FPP vs. GPP (linalool producing, Erg20p-Myc fusion) | Fig 2c, S2b | JCY51, ΔERG20::coERG20(S288C)-Myc<br>*his3::HIS3*-TEF1p_AaFSmut-Myc_SSA1t_<br>Rev(TDH3p_AaFSmut-Myc_ACT1t)<br>*leu2::LEU2*-CCW12p_SKIK-McLIS(E343D-E352H)-Myc-PDZ$_P$_ENO1t_<br>Rev(HHF2p_SKIK-McLIS(E343D-E352H)-HA-PDZ$_P$_ADH1t) | This study |

| | | | | |
|---|---|---|---|---|
| yKX728 | FPP vs. GPP (linalool producing, Erg20p-HA fusion) | S2b | JCY51, ΔERG20::coERG20(S288C)-HA<br>*his3::HIS3*-TEF1p_AaFSmut-Myc_SSA1t_<br>Rev(TDH3p_AaFSmut-Myc_ACT1t)<br>*leu2::LEU2*-<br>CCW12p_SKIK-McLIS(E343D-E352H)-Myc-<br>PDZ$_{P}$_ENO1t_<br>Rev(HHF2p_SKIK-McLIS(E343D-E352H)-<br>HA-PDZ$_{P}$_ADH1t) | This study |
| yKX607 | FPP vs. GPP (linalool producing, Erg20p-13 aa linker) | Fig 2c, S3b | JCY51, ΔERG20::coERG20(S288C)-linker1<br>*his3::HIS3*-TEF1p_AaFSmut-Myc_SSA1t_<br>Rev(TDH3p_AaFSmut-Myc_ACT1t)<br>*leu2::LEU2*-<br>CCW12p_SKIK-McLIS(E343D-E352H)-Myc-<br>PDZ$_{P}$_ENO1t_<br>Rev(HHF2p_SKIK-McLIS(E343D-E352H)-<br>HA-PDZ$_{P}$_ADH1t) | This study |
| yKX798 | FPP vs. GPP (linalool producing, 13 aa linker-Erg20p) | S3b | JCY51, ΔERG20::linker1-coERG20(S288C)<br>*his3::HIS3*-TEF1p_AaFSmut-Myc_SSA1t_<br>Rev(TDH3p_AaFSmut-Myc_ACT1t)<br>*leu2::LEU2*-<br>CCW12p_SKIK-McLIS(E343D-E352H)-Myc-<br>PDZ$_{P}$_ENO1t_<br>Rev(HHF2p_SKIK-McLIS(E343D-E352H)-<br>HA-PDZ$_{P}$_ADH1t) | This study |
| yKX795 | FPP vs. GPP (linalool producing, Erg20p-7 aa linker) | Fig 2c | JCY51, ΔERG20::coERG20(S288C)-7aa linker1<br>*his3::HIS3*-TEF1p_AaFSmut-Myc_SSA1t_<br>Rev(TDH3p_AaFSmut-Myc_ACT1t)<br>*leu2::LEU2*-<br>CCW12p_SKIK-McLIS(E343D-E352H)-Myc-<br>PDZ$_{P}$_ENO1t_<br>Rev(HHF2p_SKIK-McLIS(E343D-E352H)-<br>HA-PDZ$_{P}$_ADH1t) | This study |
| yKX796 | FPP vs. GPP (linalool producing, Erg20p-2 aa linker LE) | Fig 2c | JCY51, ΔERG20::coERG20(S288C)-2aa linker1<br>*his3::HIS3*-TEF1p_AaFSmut-Myc_SSA1t_<br>Rev(TDH3p_AaFSmut-Myc_ACT1t)<br>*leu2::LEU2*-<br>CCW12p_SKIK-McLIS(E343D-E352H)-Myc-<br>PDZ$_{P}$_ENO1t_<br>Rev(HHF2p_SKIK-McLIS(E343D-E352H)-<br>HA-PDZ$_{P}$_ADH1t) | This study |
| yKX600 | Recessive Erg20-Myc (geraniol producing, wild-type Erg20p at endogenous locus, no overexpression of Erg20p-Myc) | Fig 2g | JCY51, ΔERG20::coERG20(S288C)<br>*leu2::LEU2*-CCW12p_ObGES-Myc-<br>PDZ$_{P}$_ENO1t_<br>Rev(HHF2p_ObGES-HA-PDZ$_{P}$_ADH1t) | This study |
| yKX707 | Recessive Erg20-Myc (geraniol producing, | Fig 2g | JCY51, ΔERG20::coERG20(S288C)<br>*leu2::LEU2*-PGK1p_coERG20(S288C)- | This study |

| | | | | |
|---|---|---|---|---|
| | wild-type Erg20p at endogenous locus, overexpression of Erg20p-Myc) | | Myc_ACT1t_ CCW12p_ObGES-Myc-PDZ$_P$_ENO1t_ Rev(HHF2p_ObGES-HA-PDZ$_P$_ADH1t) | |
| yKX601 | Recessive Erg20-Myc (geraniol producing, Erg20p-Myc at endogenous locus, no overexpression of Erg20p-Myc) | Fig 2g | JCY51, ΔERG20::coERG20(S288C)-Myc *leu2::LEU2*-CCW12p_ObGES-Myc-PDZ$_P$_ENO1t_ Rev(HHF2p_ObGES-HA-PDZ$_P$_ADH1t) | This study |
| yKX778 | Recessive Erg20-Myc (geraniol producing, wild-type Erg20p at endogenous locus, overexpression of Erg20p-Myc) | Fig 2g | JCY51, ΔERG20::coERG20(S288C)-Myc *leu2::LEU2*-PGK1p_coERG20(S288C)-Myc_ACT1t_ CCW12p_ObGES-Myc-PDZ$_P$_ENO1t_ Rev(HHF2p_ObGES-HA-PDZ$_P$_ADH1t) | This study |
| yKX658 | Recessive Erg20-Myc (linalool producing, wild-type Erg20p at endogenous locus, no overexpression of Erg20p-Myc) | Fig 2h | JCY51, ΔERG20::coERG20(S288C) *leu2::LEU2*-CCW12p_SKIK-McLIS(E343D-E352H)-Myc-PDZ$_P$_ENO1t_ Rev(HHF2p_SKIK-McLIS(E343D-E352H)-HA-PDZ$_P$_ADH1t) | This study |
| yKX781 | Recessive Erg20-Myc (linalool producing, wild-type Erg20p at endogenous locus, overexpression of Erg20p-Myc) | Fig 2h | JCY51, ΔERG20::coERG20(S288C) *leu2::LEU2*-PGK1p_coERG20(S288C)-Myc_ACT1t_ CCW12p_SKIK-McLIS(E343D-E352H)-Myc-PDZ$_P$_ENO1t_ Rev(HHF2p_SKIK-McLIS(E343D-E352H)-HA-PDZ$_P$_ADH1t) | This study |
| yKX610 | Recessive Erg20-Myc (linalool producing, Erg20p-Myc at endogenous locus, no overexpression of Erg20p-Myc) | Fig 2h | JCY51, ΔERG20::coERG20(S288C)-Myc *leu2::LEU2*-CCW12p_SKIK-McLIS(E343D-E352H)-Myc-PDZ$_P$_ENO1t_ Rev(HHF2p_SKIK-McLIS(E343D-E352H)-HA-PDZ$_P$_ADH1t) | This study |
| yKX782 | Recessive Erg20-Myc (linalool producing, Erg20p-Myc at endogenous locus, no overexpression of Erg20p-Myc) | Fig 2h | JCY51, ΔERG20::coERG20(S288C)-Myc *leu2::LEU2*-PGK1p_coERG20(S288C)-Myc_ACT1t_ CCW12p_SKIK-McLIS(E343D-E352H)-Myc-PDZ$_P$_ENO1t_ Rev(HHF2p_SKIK-McLIS(E343D-E352H)-HA-PDZ$_P$_ADH1t) | This study |
| yKX813 | FPP vs. GPP (geraniol producing, Erg20p-2 aa linker fusion LK) | Fig 4d | JCY51, ΔERG20::coERG20(S288C)-LK-linker1_*his3::HIS3*-TEF1p_AaFSmut-Myc_SSA1t_ Rev(TDH3p_AaFSmut-Myc_ACT1t) *leu2::LEU2*-CCW12p_ObGES-Myc-PDZ$_P$_ENO1t_ Rev(HHF2p_ObGES-HA-PDZ$_P$_ADH1t) | This study |

| | | | | |
|---|---|---|---|---|
| yKX821 | FPP vs. GPP (geraniol producing, Erg20p-2 aa linker fusion LD) | Fig 4d | JCY51, ΔERG20::coERG20(S288C)-LD-<br>linker1_*his3::HIS3*-TEF1p_AaFSmut-<br>Myc_SSA1t_<br>Rev(TDH3p_AaFSmut-Myc_ACT1t)<br>*leu2::LEU2*-CCW12p_ObGES-Myc-<br>PDZ$_P$_ENO1t_<br>Rev(HHF2p_ObGES-HA-PDZ$_P$_ADH1t) | This study |
| yKX822 | FPP vs. GPP (geraniol producing, Erg20p-2 aa linker fusion EE) | Fig 4d | JCY51, ΔERG20::coERG20(S288C)-EE-<br>linker1_*his3::HIS3*-TEF1p_AaFSmut-<br>Myc_SSA1t_<br>Rev(TDH3p_AaFSmut-Myc_ACT1t)<br>*leu2::LEU2*-CCW12p_ObGES-Myc-<br>PDZ$_P$_ENO1t_<br>Rev(HHF2p_ObGES-HA-PDZ$_P$_ADH1t) | This study |
| yKX810 | FPP vs. GPP (geraniol producing, Erg20p-1 aa linker fusion L) | Fig 4d | JCY51, ΔERG20::coERG20(S288C)-L-<br>linker1_*his3::HIS3*-TEF1p_AaFSmut-<br>Myc_SSA1t_<br>Rev(TDH3p_AaFSmut-Myc_ACT1t)<br>*leu2::LEU2*-CCW12p_ObGES-Myc-<br>PDZ$_P$_ENO1t_<br>Rev(HHF2p_ObGES-HA-PDZ$_P$_ADH1t) | This study |
| yKX811 | FPP vs. GPP (geraniol producing, Erg20p-1 aa linker fusion E) | Fig 4d | JCY51, ΔERG20::coERG20(S288C)-E-<br>linker1_*his3::HIS3*-TEF1p_AaFSmut-<br>Myc_SSA1t_<br>Rev(TDH3p_AaFSmut-Myc_ACT1t)<br>*leu2::LEU2*-CCW12p_ObGES-Myc-<br>PDZ$_P$_ENO1t_<br>Rev(HHF2p_ObGES-HA-PDZ$_P$_ADH1t) | This study |
| yKX820 | FPP vs. GPP (geraniol producing, Erg20p-2 aa linker fusion LL) | Fig 4d | JCY51, ΔERG20::coERG20(S288C)-LL-<br>linker1_*his3::HIS3*-TEF1p_AaFSmut-<br>Myc_SSA1t_<br>Rev(TDH3p_AaFSmut-Myc_ACT1t)<br>*leu2::LEU2*-CCW12p_ObGES-Myc-<br>PDZ$_P$_ENO1t_<br>Rev(HHF2p_ObGES-HA-PDZ$_P$_ADH1t) | This study |
| yKX806 | Optimize geraniol and geranyl acetate production (2 copies of ObGES) | Fig 5a | JCY51, ΔERG20::coERG20(S288C)-Myc<br>*leu2::LEU2*-CCW12p_ObGES-Myc-<br>PDZ$_P$_ENO1t_<br>Rev(HHF2p_ObGES-HA-PDZ$_P$_ADH1t)<br>*his3::HIS3*<br>*trp1::TRP1* | This study |
| yKX807 | Optimize geraniol and geranyl acetate production (4 copies of ObGES) | Fig 5a | JCY51, ΔERG20::coERG20(S288C)-Myc<br>*leu2::LEU2*-CCW12p_ObGES-Myc-<br>PDZ$_P$_ENO1t_<br>Rev(HHF2p_ObGES-HA-PDZ$_P$_ADH1t)<br>*his3::HIS3*-CCW12p_ObGES-Myc-<br>PDZ$_P$_ENO1t_<br>Rev(PGK1p_ObGES-Myc-PDZ$_P$_ACT1t)<br>*trp1::TRP1* | This study |

| | | | | |
|---|---|---|---|---|
| yKX760 | Optimize geraniol and geranyl acetate production (6 copies of ObGES) | Fig 5a | JCY51, ΔERG20::coERG20(S288C)-Myc<br>*leu2::LEU2*-CCW12p_ObGES-Myc-PDZ$_P$_ENO1t_ Rev(HHF2p_ObGES-HA-PDZ$_P$_ADH1t)<br>*his3::HIS3*-CCW12p_ObGES-Myc-PDZ$_P$_ENO1t_ Rev(PGK1p_ObGES-Myc-PDZ$_P$_ACT1t)<br>*XII-2::KanMX*-TEF1p_ObGES-Myc-PDZ$_P$_PGK1t_ Rev(PGK1p_ObGES-Myc-PDZ$_P$_ACT1t)<br>*trp1::TRP1* | This study |
| yKX642 | Optimize geraniol and geranyl acetate production (6 copies of ObGES, overexpression of lower mevalonate pathway) | Fig 5a | JCY51, ΔERG20::coERG20(S288C)-Myc<br>*leu2::LEU2*-CCW12p_ObGES-Myc-PDZ$_P$_ENO1t_ Rev(HHF2p_ObGES-HA-PDZ$_P$_ADH1t)<br>*his3::HIS3*-CCW12p_ObGES-Myc-PDZ$_P$_ENO1t_ Rev(PGK1p_ObGES-Myc-PDZ$_P$_ACT1t)<br>*XII-2::KanMX*-TEF1p_ObGES-Myc-PDZ$_P$_PGK1t_ Rev(PGK1p_ObGES-Myc-PDZ$_P$_ACT1t)<br>*trp1::TRP1*-TDH3p_ERG19_CYC1t_TEF1p_ERG8_ACT1t_PGK1p_ERG12_ADH1t_CCW12p_IDI1_ENO1t | This study |
| yKX643 | Optimize geraniol and geranyl acetate production (6 copies of ObGES, overexpression of lower mevalonate pathway, overexpression of RhAAT) | Fig 5a | JCY51, ΔERG20::coERG20(S288C)-Myc<br>*leu2::LEU2*-CCW12p_ObGES-Myc-PDZ$_P$_ENO1t_ Rev(TDH3p_RhAAT-Myc_PGK1t)_ Rev(HHF2p_ObGES-HA-PDZ$_P$_ADH1t)<br>*his3::HIS3*-CCW12p_ObGES-Myc-PDZ$_P$_ENO1t_ Rev(PGK1p_ObGES-Myc-PDZ$_P$_ACT1t)<br>*XII-2::KanMX*-TEF1p_ObGES-Myc-PDZ$_P$_PGK1t_ Rev(PGK1p_ObGES-Myc-PDZ$_P$_ACT1t)<br>*trp1::TRP1*-TDH3p_ERG19_CYC1t_TEF1p_ERG8_ACT1t_PGK1p_ERG12_ADH1t_CCW12p_IDI1_ENO1t | This study |
| yKX809 | Optimize linalool and linalyl acetate production (2 copies of ObGES) | Fig 5b | JCY51, ΔERG20::coERG20(S288C)-Myc<br>*leu2::LEU2*-CCW12p_SKIK-McLIS(E343D-E352H)-Myc-PDZ$_P$_ENO1t_ Rev(HHF2p_SKIK-McLIS(E343D-E352H)-HA-PDZ$_P$_ADH1t)<br>*his3::HIS3*<br>*trp1::TRP1* | This study |

| | | | | |
|---|---|---|---|---|
| yKX808 | Optimize linalool and linalyl acetate production (4 copies of ObGES) | Fig 5b | JCY51, ΔERG20::coERG20(S288C)-Myc<br>*leu2::LEU2*-<br>CCW12p_SKIK-McLIS(E343D-E352H)-Myc-PDZ$_P$_ENO1t_<br>Rev(HHF2p_SKIK-McLIS(E343D-E352H)-HA-PDZ$_P$_ADH1t)<br>*his3::HIS3*-<br>CCW12p_SKIK-McLIS(E343D-E352H)-Myc-PDZ$_P$_ENO1t_<br>Rev(TDH3p_SKIK-McLIS(E343D-E352H)-HA-PDZ$_P$_PGK1t)<br>*trp1::TRP1* | This study |
| yKX803 | Optimize linalool and linalyl acetate production (6 copies of ObGES) | Fig 5b | JCY51, ΔERG20::coERG20(S288C)-Myc<br>*leu2::LEU2*-<br>CCW12p_SKIK-McLIS(E343D-E352H)-Myc-PDZ$_P$_ENO1t_<br>Rev(HHF2p_SKIK-McLIS(E343D-E352H)-HA-PDZ$_P$_ADH1t)<br>*his3::HIS3*-<br>CCW12p_SKIK-McLIS(E343D-E352H)-Myc-PDZ$_P$_ENO1t_<br>Rev(TDH3p_SKIK-McLIS(E343D-E352H)-HA-PDZ$_P$_PGK1t)<br>*XII-2::KanMX*-<br>CCW12p_SKIK-McLIS(E343D-E352H)-Myc-PDZ$_P$_ENO1t_<br>Rev(PGK1p_SKIK-McLIS(E343D-E352H)-HA-PDZ$_P$_ACT1t)<br>*trp1::TRP1* | This study |
| yKX734 | Optimize linalool and linalyl acetate production (6 copies of ObGES, overexpression of lower mevalonate pathway) | Fig 5b | JCY51, ΔERG20::coERG20(S288C)-Myc<br>*leu2::LEU2*-<br>CCW12p_SKIK-McLIS(E343D-E352H)-Myc-PDZ$_P$_ENO1t_<br>Rev(HHF2p_SKIK-McLIS(E343D-E352H)-HA-PDZ$_P$_ADH1t)<br>*his3::HIS3*-<br>CCW12p_SKIK-McLIS(E343D-E352H)-Myc-PDZ$_P$_ENO1t_<br>Rev(TDH3p_SKIK-McLIS(E343D-E352H)-HA-PDZ$_P$_PGK1t)<br>*XII-2::KanMX*-<br>CCW12p_SKIK-McLIS(E343D-E352H)-Myc-PDZ$_P$_ENO1t_<br>Rev(PGK1p_SKIK-McLIS(E343D-E352H)-HA-PDZ$_P$_ACT1t)<br>*trp1::TRP1*-<br>TDH3p_ERG19_CYC1t_TEF1p_ERG8_ACT1t<br>_<br>PGK1p_ERG12_ADH1t_CCW12p_IDI1_ENO<br>1t | This study |
| yKX735 | Optimize linalool and linalyl acetate | Fig 5b | JCY51, ΔERG20::coERG20(S288C)-Myc<br>*leu2::LEU2*-TEF1p_CbAAT9-1-c- | This study |

| | | | | |
|---|---|---|---|---|
| | production (6 copies of ObGES, overexpression of lower mevalonate pathway, 2 copies of CbAAT) | | Myc_SSA1t_<br>CCW12p_SKIK-McLIS(E343D-E352H)-Myc-PDZ$_P$_ENO1t_<br>Rev(TDH3p_CbAAT9-1-c-Myc_PGK1t)_<br>Rev(HHF2p_SKIK-McLIS(E343D-E352H)-HA-PDZ$_P$_ADH1t)<br>*his3::HIS3*-<br>CCW12p_SKIK-McLIS(E343D-E352H)-Myc-PDZ$_P$_ENO1t_<br>Rev(TDH3p_SKIK-McLIS(E343D-E352H)-HA-PDZ$_P$_PGK1t)<br>*XII-2::KanMX*-<br>CCW12p_SKIK-McLIS(E343D-E352H)-Myc-PDZ$_P$_ENO1t_<br>Rev(PGK1p_SKIK-McLIS(E343D-E352H)-HA-PDZ$_P$_ACT1t)<br>*trp1::TRP1*-<br>TDH3p_ERG19_CYC1t_TEF1p_ERG8_ACT1t_<br>PGK1p_ERG12_ADH1t_CCW12p_IDI1_ENO1t | |
| yKX736 | Optimize linalool and linalyl acetate production (6 copies of ObGES, overexpression of lower mevalonate pathway, 4 copies of CbAAT) | Fig 5b | JCY51, ΔERG20::coERG20(S288C)-Myc<br>*leu2::LEU2*-TEF1p_CbAAT9-1-c-Myc_SSA1t_<br>CCW12p_SKIK-McLIS(E343D-E352H)-Myc-PDZ$_P$_ENO1t_<br>Rev(TDH3p_CbAAT9-1-c-Myc_PGK1t)_<br>Rev(HHF2p_SKIK-McLIS(E343D-E352H)-HA-PDZ$_P$_ADH1t)<br>*his3::HIS3*-TDH3p_CbAAT9-1-c-Myc_PGK1t<br>CCW12p_SKIK-McLIS(E343D-E352H)-Myc-PDZ$_P$_ENO1t_<br>Rev(TEF1p_CbAAT9-1-c-Myc_SSA1t)<br>Rev(TDH3p_SKIK-McLIS(E343D-E352H)-HA-PDZ$_P$_PGK1t)<br>*XII-2::KanMX*-<br>CCW12p_SKIK-McLIS(E343D-E352H)-Myc-PDZ$_P$_ENO1t_<br>Rev(PGK1p_SKIK-McLIS(E343D-E352H)-HA-PDZ$_P$_ACT1t)<br>*trp1::TRP1*-<br>TDH3p_ERG19_CYC1t_TEF1p_ERG8_ACT1t_<br>PGK1p_ERG12_ADH1t_CCW12p_IDI1_ENO1t | This study |
| yKX737 | Optimize linalool and linalyl acetate production (6 copies of ObGES, overexpression of lower mevalonate pathway, 6 copies of CbAAT) | Fig 5b | JCY51, ΔERG20::coERG20(S288C)-Myc<br>*leu2::LEU2*-TEF1p_CbAAT9-1-c-Myc_SSA1t_<br>CCW12p_SKIK-McLIS(E343D-E352H)-Myc-PDZ$_P$_ENO1t_<br>Rev(TDH3p_CbAAT9-1-c-Myc_PGK1t)_<br>Rev(HHF2p_SKIK-McLIS(E343D-E352H)-HA-PDZ$_P$_ADH1t) | This study |

| | | | | |
|---|---|---|---|---|
| | | | *his3::HIS3*-TDH3p_CbAAT9-1-c-Myc_PGK1t CCW12p_SKIK-McLIS(E343D-E352H)-Myc-PDZ$_P$_ENO1t_ Rev(TEF1p_CbAAT9-1-c-Myc_SSA1t) Rev(TDH3p_SKIK-McLIS(E343D-E352H)-HA-PDZ$_P$_PGK1t)<br>*XII-2::KanMX*-TDH3p_CbAAT9-1-c-Myc_PGK1t CCW12p_SKIK-McLIS(E343D-E352H)-Myc-PDZ$_P$_ENO1t_ Rev(TEF1p_CbAAT9-1-c-Myc_SSA1t) Rev(PGK1p_SKIK-McLIS(E343D-E352H)-HA-PDZ$_P$_ACT1t)<br>*trp1::TRP1*-TDH3p_ERG19_CYC1t_TEF1p_ERG8_ACT1t_ PGK1p_ERG12_ADH1t_CCW12p_IDI1_ENO1t | |
| yKX638 | Recruitment of Erg20p-Myc to condensate (farnesene producing) | Fig 6b | JCY51, ΔERG20::coERG20(S288C)-Myc-PDZ$_P$<br>*leu2::LEU2*-TEF1p_ aaFSmut-Myc_SSA1t_ Rev(TDH3p_aaFSmut-Myc_ACT1t)<br>*his3::HIS3*-Rev(TDH3p_PDZ-PDZ-FUSn-mCherry-I301(K129A)_TDH1t)_TEF1p_PDZ-PDZ-FUSn-mCherry-I301(K129A)_TDH1t | This study |
| yKX639 | No recruitment of Erg20p to condensate (farnesene producing) | Fig 6b | JCY51, ΔERG20::coERG20(S288C)-Myc-PDZ$_P$<br>*leu2::LEU2*-TEF1p_ aaFSmut-Myc_SSA1t_ Rev(TDH3p_aaFSmut-Myc_ACT1t)<br>*his3::HIS3*-Rev(TDH3p_FUSn -mCherry-I301(K129A)_TDH1t)<br>TEF1p_FUSn-mCherry-I301(K129A)_TDH1t | This study |
| yKX640 | Recruitment of Erg20p(F96W) to condensate (farnesene producing) | Fig 6b | JCY51, ΔERG20::coERG20(F96W)(S288C)-Myc-PDZ$_P$<br>*leu2::LEU2*-TEF1p_ aaFSmut-Myc_SSA1t_ Rev(TDH3p_aaFSmut-Myc_ACT1t)<br>*his3::HIS3*-Rev(TDH3p_PDZ-PDZ-FUSn-mCherry-I301(K129A)_TDH1t)_TEF1p_PDZ-PDZ-FUSn-mCherry-I301(K129A)_TDH1t | This study |
| yKX641 | No recruitment of Erg20p(F96W) to condensate (farnesene producing) | Fig 6b | JCY51, ΔERG20::coERG20(F96W)(S288C)-Myc-PDZ$_P$<br>*leu2::LEU2*-TEF1p_ aaFSmut-Myc_SSA1t_ Rev(TDH3p_aaFSmut-Myc_ACT1t)<br>*his3::HIS3*-Rev(TDH3p-FUSn-mCherry- | This study |

| | | | | |
|---|---|---|---|---|
| | | | I301(K129A)_TDH1t)_TEF1p_ FUSn-mCherry-I301(K129A)_TDH1t | |
| yKX710 | Colocalize Erg20p and ObGES (geraniol producing, wildtype Erg20p-Myc recruited with ObGES) | Fig 6d, S7a, S7b | JCY51, ΔERG20::coERG20(S288C)-Myc-PDZ$_P$ *leu2::LEU2*-CCW12p_ObGES-Myc-PDZ$_P$_ENO1t_ TDH3p_PDZ-PDZ-FUSn-mCherry-I301(K129A)_ADH1t_ Rev(TEF1p_PDZ-PDZ-FUSn-mCherry-I301(K129A)_TDH1t)_ Rev(HHF2p_ObGES-HA-PDZ$_P$_ADH1t) *his3::HIS3*-Rev(TDH3p_PDZ-PDZ-FUSn-mCherry-I301(K129A)_ADH1t)_ CCW12p_ObGES-Myc-PDZ$_P$_ENO1t_ TEF1p_PDZ-PDZ-FUSn-mCherry-I301(K129A)_TDH1t_ Rev(PGK1p_ObGES-Myc-PDZ$_P$_ACT1t) *XII-2::KanMX*-TEF1p_ObGES-Myc-PDZ$_P$_PGK1t_ CCW12p_PDZ-PDZ-FUSn-mCherry-I301(K129A)_ENO1t_ Rev(PGK1p_ObGES-Myc-PDZ$_P$_ACT1t)_ Rev(TDH3p_PDZ-PDZ-FUSn-mCherry-I301(K129A)_TDH1t) *trp1::TRP1*-TDH3p_ERG19_CYC1t_TEF1p_ERG8_ACT1t_ PGK1p_ERG12_ADH1t_CCW12p_IDI1_ENO1t | This study |
| yKX711 | Colocalize Erg20p and ObGES (geraniol producing, wildtype Erg20p-Myc and ObGES not recruited) | Fig 6d, S7a, S7b | JCY51, ΔERG20::coERG20(S288C)-Myc-PDZ$_P$ *leu2::LEU2*-CCW12p_ObGES-Myc-PDZ$_P$_ENO1t_ TDH3p_FUSn-mCherry-I301(K129A)_ADH1t_ Rev(TEF1p_FUSn-mCherry-I301(K129A)_TDH1t)_ Rev(HHF2p_ObGES-HA-PDZ$_P$_ADH1t) *his3::HIS3*-Rev(TDH3p_FUSn-mCherry-I301(K129A)_ADH1t)_ CCW12p_ObGES-Myc-PDZ$_P$_ENO1t_ TEF1p_ FUSn-mCherry-I301(K129A)_TDH1t_ Rev(PGK1p_ObGES-Myc-PDZ$_P$_ACT1t) *XII-2::KanMX*-TEF1p_ObGES-Myc-PDZ$_P$_PGK1t_ CCW12p_FUSn-mCherry-I301(K129A)_ENO1t_ Rev(PGK1p_ObGES-Myc-PDZ$_P$_ACT1t)_ Rev(TDH3p_FUSn-mCherry-I301(K129A)_TDH1t) *trp1::TRP1*-TDH3p_ERG19_CYC1t_TEF1p_ERG8_ACT1t_ | This study |

| | | | | |
|---|---|---|---|---|
| | | | PGK1p_ERG12_ADH1t_CCW12p_IDI1_ENO1t | |
| yKX712 | Colocalize Erg20p and ObGES (Erg20p(F96W)-Myc recruited with ObGES) | Fig 6d, S8b | JCY51, ΔERG20::coERG20(F96W)(S288C)-Myc-PDZ$_P$ *leu2::LEU2*-CCW12p_ObGES-Myc-PDZ$_P$_ENO1t_ TDH3p_PDZ-PDZ-FUSn-mCherry-I301(K129A)_ADH1t_ Rev(TEF1p_PDZ-PDZ-FUSn-mCherry-I301(K129A)_TDH1t)_ Rev(HHF2p_ObGES-HA-PDZ$_P$_ADH1t) *his3::HIS3*-Rev(TDH3p_PDZ-PDZ-FUSn-mCherry-I301(K129A)_ADH1t)_ CCW12p_ObGES-Myc-PDZ$_P$_ENO1t_ TEF1p_PDZ-PDZ-FUSn-mCherry-I301(K129A)_TDH1t_ Rev(PGK1p_ObGES-Myc-PDZ$_P$_ACT1t) *XII-2::KanMX*-TEF1p_ObGES-Myc-PDZ$_P$_PGK1t_ CCW12p_PDZ-PDZ-FUSn-mCherry-I301(K129A)_ENO1t_ Rev(PGK1p_ObGES-Myc-PDZ$_P$_ACT1t)_ Rev(TDH3p_PDZ-PDZ-FUSn-mCherry-I301(K129A)_TDH1t) *trp1::TRP1*-TDH3p_ERG19_CYC1t_TEF1p_ERG8_ACT1t_ PGK1p_ERG12_ADH1t_CCW12p_IDI1_ENO1t | This study |
| yKX713 | Colocalize Erg20p and ObGES (geraniol producing, Erg20p(F96W)-Myc and ObGES not recruited to condensate) | Fig 6d, S8b | JCY51, ΔERG20::coERG20(F96W)(S288C)-Myc-PDZ$_P$ *leu2::LEU2*-CCW12p_ObGES-Myc-PDZ$_P$_ENO1t_ TDH3p_FUSn-mCherry-I301(K129A)_ADH1t_ Rev(TEF1p_FUSn-mCherry-I301(K129A)_TDH1t)_ Rev(HHF2p_ObGES-HA-PDZ$_P$_ADH1t) *his3::HIS3*-Rev(TDH3p_FUSn-mCherry-I301(K129A)_ADH1t)_ CCW12p_ObGES-Myc-PDZ$_P$_ENO1t_ TEF1p_ FUSn-mCherry-I301(K129A)_TDH1t_ Rev(PGK1p_ObGES-Myc-PDZ$_P$_ACT1t) *XII-2::KanMX*-TEF1p_ObGES-Myc-PDZ$_P$_PGK1t_ CCW12p_FUSn-mCherry-I301(K129A)_ENO1t_ Rev(PGK1p_ObGES-Myc-PDZ$_P$_ACT1t)_ Rev(TDH3p_FUSn-mCherry-I301(K129A)_TDH1t) *trp1::TRP1*-TDH3p_ERG19_CYC1t_TEF1p_ERG8_ACT1t_ | This study |

| | | | | |
|---|---|---|---|---|
| | | | PGK1p_ERG12_ADH1t_CCW12p_IDI1_ENO1t | |
| yKX714 | Colocalize Erg20p and ObGES (geranyl acetate producing, Erg20p-Myc recruited with ObGES) | S8b | JCY51, ΔERG20::coERG20(S288C)-Myc-PDZ$_P$ *leu2::LEU2*-CCW12p_ObGES-Myc-PDZ$_P$_ENO1t_ TDH3p_PDZ-PDZ-FUSn-mCherry-I301(K129A)_ADH1t_ Rev(TEF1p_PDZ-PDZ-FUSn-mCherry-I301(K129A)_TDH1t)_ Rev(TDH3p_RhAAT-Myc_PGK1t)_ Rev(HHF2p_ObGES-HA-PDZ$_P$_ADH1t) *his3::HIS3*-Rev(TDH3p_PDZ-PDZ-FUSn-mCherry-I301(K129A)_ADH1t)_ CCW12p_ObGES-Myc-PDZ$_P$_ENO1t_ TEF1p_PDZ-PDZ-FUSn-mCherry-I301(K129A)_TDH1t_ Rev(PGK1p_ObGES-Myc-PDZ$_P$_ACT1t) *XII-2::KanMX*-TEF1p_ObGES-Myc-PDZ$_P$_PGK1t_ CCW12p_PDZ-PDZ-FUSn-mCherry-I301(K129A)_ENO1t_ Rev(PGK1p_ObGES-Myc-PDZ$_P$_ACT1t)_ Rev(TDH3p_PDZ-PDZ-FUSn-mCherry-I301(K129A)_TDH1t) *trp1::TRP1*-TDH3p_ERG19_CYC1t_TEF1p_ERG8_ACT1t_ PGK1p_ERG12_ADH1t_CCW12p_IDI1_ENO1t | This study |
| yKX715 | Colocalize Erg20p and ObGES (geranyl acetate producing, Erg20p-Myc and ObGES not recruited) | S8b | JCY51, ΔERG20::coERG20(S288C)-Myc-PDZ$_P$ *leu2::LEU2*-CCW12p_ObGES-Myc-PDZ$_P$_ENO1t_ TDH3p_FUSn-mCherry-I301(K129A)_ADH1t_ Rev(TEF1p_FUSn-mCherry-I301(K129A)_TDH1t)_ Rev(TDH3p_RhAAT-Myc_PGK1t)_ Rev(HHF2p_ObGES-HA-PDZ$_P$_ADH1t) *his3::HIS3*-Rev(TDH3p_FUSn-mCherry-I301(K129A)_ADH1t)_ CCW12p_ObGES-Myc-PDZ$_P$_ENO1t_ TEF1p_ FUSn-mCherry-I301(K129A)_TDH1t_ Rev(PGK1p_ObGES-Myc-PDZ$_P$_ACT1t) *XII-2::KanMX*-TEF1p_ObGES-Myc-PDZ$_P$_PGK1t_ CCW12p_FUSn-mCherry-I301(K129A)_ENO1t_ Rev(PGK1p_ObGES-Myc-PDZ$_P$_ACT1t)_ Rev(TDH3p_FUSn-mCherry-I301(K129A)_TDH1t) *trp1::TRP1*- | This study |

| | | | | |
|---|---|---|---|---|
| | | | TDH3p_ERG19_CYC1t_TEF1p_ERG8_ACT1t_PGK1p_ERG12_ADH1t_CCW12p_IDI1_ENO1t | |
| yKX716 | Colocalize Erg20p and ObGES (geranyl acetate producing, Erg20p(F96W)-Myc and ObGES recruited) | S8b | JCY51, ΔERG20::coERG20(F96W)(S288C)-Myc-PDZ$_P$ *leu2::LEU2*-CCW12p_ObGES-Myc-PDZ$_P$_ENO1t_TDH3p_PDZ-PDZ-FUSn-mCherry-I301(K129A)_ADH1t_Rev(TEF1p_PDZ-PDZ-FUSn-mCherry-I301(K129A)_TDH1t)_Rev(TDH3p_RhAAT-Myc_PGK1t)_Rev(HHF2p_ObGES-HA-PDZ$_P$_ADH1t) *his3::HIS3*-Rev(TDH3p_PDZ-PDZ-FUSn-mCherry-I301(K129A)_ADH1t)_CCW12p_ObGES-Myc-PDZ$_P$_ENO1t_TEF1p_PDZ-PDZ-FUSn-mCherry-I301(K129A)_TDH1t_ Rev(PGK1p_ObGES-Myc-PDZ$_P$_ACT1t) *XII-2::KanMX*-TEF1p_ObGES-Myc-PDZ$_P$_PGK1t_CCW12p_PDZ-PDZ-FUSn-mCherry-I301(K129A)_ENO1t_ Rev(PGK1p_ObGES-Myc-PDZ$_P$_ACT1t)_Rev(TDH3p_PDZ-PDZ-FUSn-mCherry-I301(K129A)_TDH1t) *trp1::TRP1*-TDH3p_ERG19_CYC1t_TEF1p_ERG8_ACT1t_PGK1p_ERG12_ADH1t_CCW12p_IDI1_ENO1t | This study |
| yKX717 | Colocalize Erg20p and ObGES (geranyl acetate producing, Erg20p(F96W)-Myc and ObGES not recruited) | S8b | JCY51, ΔERG20::coERG20(F96W)(S288C)-Myc-PDZ$_P$ *leu2::LEU2*-CCW12p_ObGES-Myc-PDZ$_P$_ENO1t_TDH3p_FUSn-mCherry-I301(K129A)_ADH1t_Rev(TEF1p_FUSn-mCherry-I301(K129A)_TDH1t)_Rev(TDH3p_RhAAT-Myc_PGK1t)_Rev(HHF2p_ObGES-HA-PDZ$_P$_ADH1t) *his3::HIS3*-Rev(TDH3p_FUSn-mCherry-I301(K129A)_ADH1t)_CCW12p_ObGES-Myc-PDZ$_P$_ENO1t_TEF1p_ FUSn-mCherry-I301(K129A)_TDH1t_Rev(PGK1p_ObGES-Myc-PDZ$_P$_ACT1t) *XII-2::KanMX*-TEF1p_ObGES-Myc-PDZ$_P$_PGK1t_CCW12p_FUSn-mCherry-I301(K129A)_ENO1t_ Rev(PGK1p_ObGES-Myc-PDZ$_P$_ACT1t)_Rev(TDH3p_FUSn-mCherry- | This study |

| | | | |
|---|---|---|---|
| | | | I301(K129A)_TDH1t)<br>*trp1::TRP1*-<br>TDH3p_ERG19_CYC1t_TEF1p_ERG8_ACT1_PGK1p_ERG12_ADH1t_CCW12p_IDI1_ENO1t |
| yKX786 | Variant of yKX712 for bioreactor | Fig 6f | JCY51, ΔERG20::coERG20mut(F96W)(S288C)-Myc-PDZL<br>*leu2::LEU2*-CCW12p_obGES-Myc-PDZL_ENO1t_TDH3p_PDZ-PDZ-FUSn-mCherry-I301(K129A)_ADH1t_Rev(TEF1p_PDZ-PDZ-FUSn-mCherry-I301(K129A)_TDH1t)-_Rev(HHF2p_obGES-HA-PDZL_ADH1t)<br>*his3::HIS3*-Rev(TDH3p_PDZ-PDZ-FUSn-mCherry-I301(K129A)_TDH1t)_CCW12p_obGES-Myc-PDZL_ENO1t_TEF1p_PDZ-PDZ-FUSn-mCherry-I301(K129A)_TDH1t_Rev(PGK1p_obGES-Myc-PDZL_ACT1t)_XII-2-TEF1p_obGES-Myc-PDZL_PGK1t_CCW12p_PDZ-PDZ-FUSn-mCherry-I301(K129A)_ENO1t_Rev(PGK1p_obGES-Myc-PDZL_ACT1t)_Rev(TDH3p_PDZ-PDZ-FUSn-mCherry-I301(K129A)_TDH1t)<br>*trp1::TRP1*-TDH3p_ERG19_CYC1t_TEF1p_ERG8_ACT1t_PGK1p_ERG12_ADH1t_CCW12p_IdI1(wt)_ENO1t<br>*ura3::URA3* |

**Supplementary Note 1**

We examine the relationship between GPP reaction and diffusion within a biomolecular condensate through the Damköhler number, which for a first-order reaction is given by the equation:

$$\mathrm{Da} = \frac{\mathrm{rxn}}{\mathrm{diffusion}} = \frac{kR^2}{\mathcal{D}} \qquad (1)$$

The reaction constant k can be found in Table 1. The radius of the condensate R can be measured from microscopy, as in Figure 1, where R≈0.625 * $10^{-6}$ m. The diffusion constant $\mathcal{D}$ can be found through the Stokes-Einstein relation:

$$\mathcal{D} = \frac{k_b T}{6\pi\eta r} \quad (2)$$

We assume that GPP can be modeled as a sphere of hydrodynamic radius $r \approx 0.5$ nm, based on its chemical structure. We use T = 303K and take $\eta_{FUS} \approx 1$ Pa-s.[4] With these assumptions, in the condensate dense phase, we find $Da_{Erg20p\text{-}Myc,\ GPP} \approx 4.2$ and $Da_{Erg20p(F96W)\text{-}Myc,\ GPP} \approx 1.9$. In the dilute phase, we take dynamic viscosity to be similar to that of water, and find that $Da_{Erg20p\text{-}Myc,\ GPP}$ and $Da_{Erg20p(F96W)\text{-}Myc,\ GPP} << 1$. When GPP is in FUS condensates, $Da_{GPP}$ is O(1) and so enzyme kinetics has a comparable effect as mass-transfer, while in the dilute phase, the system is mass-transfer limited. Therefore, the decrease in $k_{cat,\ GPP}$ observed in the Erg20p(F96W)-Myc relative to Erg20p-Myc *in vitro* could be implicated in the relative flux to geraniol over farnesene *in vivo*.

**Reference**


Shaw, W. M. *et al.* Engineering a Model Cell for Rational Tuning of GPCR Signaling. *Cell* **177**, 782-796.e27 (2019).

Entian, K. D. & Kötter, P. 25 Yeast Genetic Strain and Plasmid Collections. *Methods in Microbiology* **36**, 629–666 (2007).

Wegner, S. A. *et al.* Engineering acetyl-CoA supply and *ERG9* repression to enhance mevalonate production in *Saccharomyces cerevisiae*. *J Ind Microbiol Biotechnol* **48**, (2021).

Shen, Y. *et al.* The liquid-to-solid transition of FUS is promoted by the condensate surface. *Proceedings of the National Academy of Sciences*, **120** (2023).
